# High-throughput histopathology for complex in vitro models

**DOI:** 10.1101/2025.04.01.646522

**Authors:** Marius F. Harter, Elisa D’Arcangelo, Julien Aubert, Barbora Lavickova, Charles Havnar, Bilgenaz Stoll, Irineja Cubela, Adrian M. Filip, Laura Gaspa-Toneu, Julian Scherer, Evdoxia Karagkiozi, Jean-Samuel Dupre, Rupén López-Sandoval, Rebecca Hsia, Leah M. Norona, Giovanna Brancati, Ryo Okuda, J. Gray Camp, Eliah R. Shamir, Jose Luis Garcia-Cordero, Iago Pereiro, Nikolche Gjorevski

## Abstract

Human complex in vitro models (CIVMs) have demonstrated remarkable potential to study tissue development, physiology and disease at high-throughput. To effectively employ these miniaturized systems in translational preclinical research, their in-depth benchmarking is pivotal. Histology has been the core of tissue characterization for centuries and the foundation of spatial phenotyping. However, standard histology workflows are inherently low-throughput and centered on large tissue pieces. This does not match the high sample volumes and small sample sizes in CIVM research. Here, we introduce a holistic ‘histo-workflow’, utilizing 3D-printed histomolds that facilitate co-planar embedding of CIVMs at high-throughput, resulting in up to 48 samples in one section. We developed a variety of model-specific histomold designs that enable spatially controlled histological sectioning and downstream analyses. We describe these workflows, including mold generation, highplex staining and image analysis, and exemplify their application to histological analyses of various CIVMs. Altogether, the histomolds introduced here afford opportunities for CIVM processing and analysis, while significantly reducing labor and reagent resources, thereby democratizing high-throughput CIVM in histopathology.

## Introduction

CIVMs have emerged as revolutionary tools to study human physiology^1–5^. These - often patient-derived - three dimensional (3D) in vitro systems generally surpass conventional 2D models as tissue and organ mimetics, as they more accurately recapitulate the architecture and functionality of the respective organ of origin, therefore opening new avenues to understand human tissue function in health and disease^6–10^. Among the most notable models are patient-derived explants^11–13^, organ-on-chip systems^14–17^, barrier tissues grown on transwells^18–21^, as well as organoids^22–27^, either induced-pluripotent stem cell- (iPSC) or adult stem cell-derived, which can be subjected to various (culture) methods and even supplemented with immune or stromal cells to study organ-level processes^28–32^. Due to the diversity of the aforementioned models combined with a vast number of permutations in culture techniques, culture duration, culture vessels, media formulations and co-culture strategies implemented by the scientific community thus far, comprehensive, fast and reliable characterization of such lab-grown structures is critical for their effective integration into basic and translational research^33–35^.

Histology has been a cornerstone for primary organ assessment and CIVM characterization for decades, as it provides insight into the cellular composition, differentiation status, spatial context and integrity of the tissue structures examined. Critically, it also enables direct comparison of CIVMs to the parental donor tissues, which act as both a blueprint and benchmark. To assess CIVMs with histological readouts, formalin-fixed, paraffin-embedded (FFPE) blocks are typically prepared per individual sample and subsequently sectioned using a microtome, resulting in thin sections of the specimen. These sections can then be subjected to a wide range of downstream assays, such as hematoxylin and eosin staining (H&E), immunohistochemistry (IHC), multiplex immunofluorescence (mIF), special stains or spatial omics techniques to evaluate the physiological relevance of the CIVM.

Despite the wealth of information that histological readouts offer, it is traditionally a very low-throughput method, characterized by time-consuming, manual workflows that are not well-suited to accommodate large sample volumes^36,37^. These drawbacks are particularly pronounced in the case of CIVMs as samples are often very small size (e.g. 1-day old intestinal organoids ∼ 100 µm diameter), demand complex culture vessels (e.g. transwells or organ-on-chips) and culture methods (e.g. suspension vs. extracellular matrix (ECM) culture) that complicate the handling and embedding workflows significantly, while yielding low biological material per section. In addition, one key advantage of CIVMs is the possibility of extensive experimental designs that include many conditions and, critically, the opportunity for longitudinal studies with multiple time points. Altogether, this comes at the cost of insurmountable sample numbers that are incompatible with conventional histology workflows.

Recent efforts have aimed to address the aforementioned challenges of embedding organoids for histopathological analysis. Zhang et al.^38^ utilized centrifugation to concentrate organoid samples in agarose pellets, while others deployed Histogel™ to embed organoids directly from culture plates^39^. While these methods increased the organoid density per section, they did not enhance FFPE-embedding throughput, still demanding sample-by-sample processing. Inspired by tissue microarrays^40–42^, some groups have developed negative molds for high-throughput co-planar embedding of zebrafish and CIVMs^43–46^. However, these workflows require advanced fabrication methods like CNC-milling and PDMS-lithography, which are infeasible to implement for standard histology labs. Additionally, these models are not easily adaptable to the diverse landscape of emerging *in vitro* models. On top, current protocols focus on large-scale models (>800 µm), such as neural organoids, which are comparably simple to handle^36,37,46–48^. More complex models such as epithelial monolayers cultured on transwells, organoids grown in an ECM or organoids co-cultured with immune cells remain unaddressed, highlighting the need for a pragmatic solution to integrate diverse CIVMs into histopathological workflows efficiently.

Here, we present a comprehensive ‘histo-workflow’ to overcome current limitations in CIVM FFPE-processing, data acquisition and analysis. Our approach is demonstrated across various CIVM types, including organoids cultured in matrix, immune-epithelium co-cultures and transwells. This blueprint stands on two key pillars: first, 3D-printed ‘histomolds’ to enable straightforward, co-planar high-throughput embedding of a wide range of CIVMs. Second, an optimized workflow for sample transfer, FFPE-embedding and identical staining and analysis of dozens of samples within a single section. Beyond that, we provide a novel protocol for preserving spatial relationships in ECM-embedded cultures, enabling the study of spatially integer epithelial-immune cell interactions. Finally, we validate our workflow with extensive datasets using conventional FFPE-staining techniques and a feasibility study for spatial transcriptomics.

## Results

### Histo-workflow for complex in vitro systems at high-throughput

We describe a blueprint to embed CIVMs in histopathology workflows at high-throughput facilitated by ‘histomolds’. The histomolds were designed and 3D printed to address the embedding requirements of a large variety of CIVMs, ultimately accelerating phenotypic characterization of these systems within preclinical research. Most crucially, the approach described here does not require any modification of the conventional histology workflows (Fig. 1a; Suppl. Fig. 1a). Briefly, 3D-printed histomolds contain an array of protrusions of round or rectangular shapes. Their overall size is governed by the dimensions of standard biopsy-casettes (32 x 26mm), metallic molds (32 x 24mm) for FFPE-blocks and conventional glass-slides (75 x 25mm). After filling the histomolds with liquid Histogel™ and allowing the gel to polymerize, the resulting countermold comprises an array of round or rectangular wells (Fig. 1a; Suppl. Fig. 1a), enabling co-planar embedding of up to 48 samples. Hereafter, we refer to the resulting countermold as “histoarray”. Once transferred into the histoarray, the samples are covered with an additional layer of liquid Histogel™ to obtain a histoarray that encapsulates all samples (Fig. 1a, Suppl. Fig. 1b). This histoarray block is further processed using standard procedures to generate FFPE-blocks (Suppl. Fig. 1c).

**Fig. 1:**
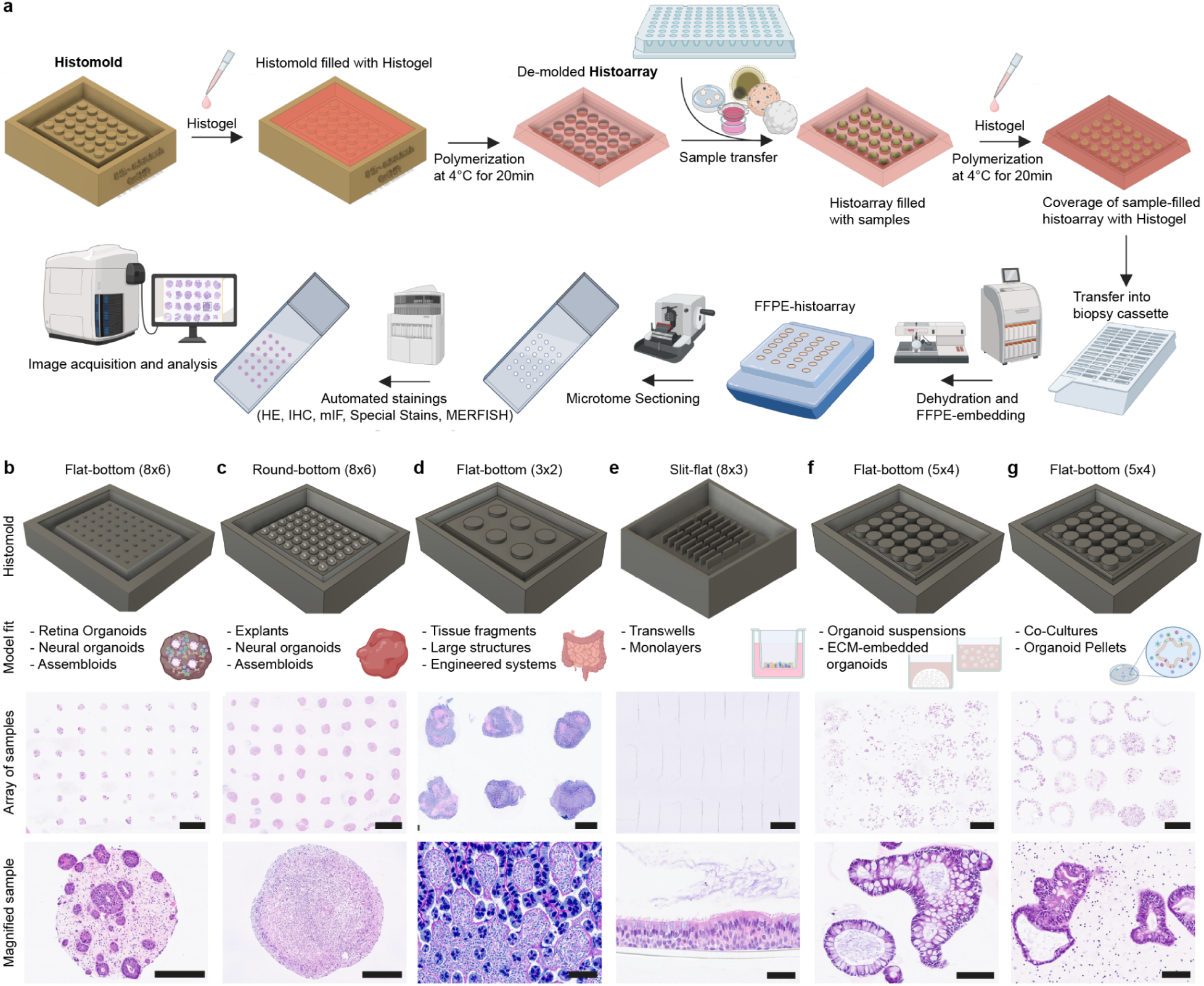
High-throughput histo-workflow facilitated by histomolds for complex in vitro systems in histopathology. **a**, Illustration of the entire histo-workflow. **b-g** Non-exhaustive histomold overview over a span of routinely used in vitro models and respective culture protocols, ordered by ease of embedding. (Histoarray overview scale bar: 2.5 mm). **b,** Forty-five single, mid-sized colorectal cancer assembloids embedded by direct transfer into histoarray. Magnified H&E highlights a single assembloid (Scale bar: 250 µm). **c,** Forty-seven single, mouse neural organoids embedded by direct transfer into histoarray. Magnified H&E highlights a single organoid (Scale bar: 250 µm). **d,** Six intestinal tissues embedded by direct transfer in histoarray. Magnified Alcian Blue staining exemplifies a region of interest in the tissue (Scale bar: 100 µm). **e,** Twenty-four alveolar epithelia cultured on 96-well transwells embedded by peeling away the specimen-polymer complex with forceps and transfer in the histoarray. Magnified H&E exemplifies ciliated, bronchial epithelium cultured on 24-well transwells with apically secreted mucus (refer to Fig. 4; Scale bar: 50 µm). **f,** Histoarray containing twenty wells of soft ECM-cultured (e.g. Matrigel) intestinal organoids (refer to Fig. 5; Scale bar: 2 mm). Magnified H&E represents two intestinal organoids out of a dozen within the same sample-well (Scale bar: 100 µm). **g,** Histoarray of twenty stiff ECM-embedded (e.g. Collagen-Matrigel) PBMC-intestinal organoid co-cultures (refer to Fig. 6; Scale bar: 100 µm).

**Suppl. Fig. 1:**
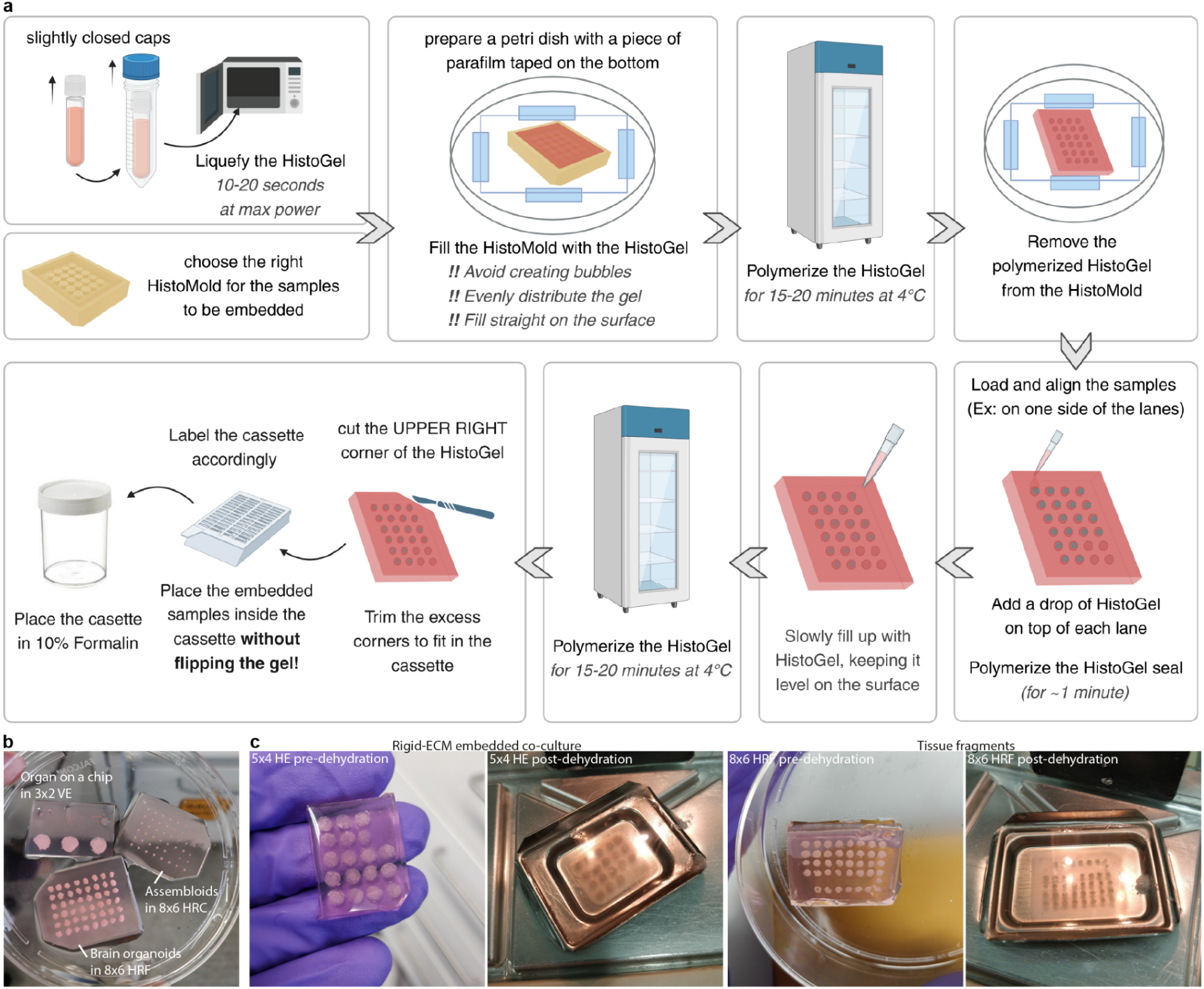
Histoarray generation and preparation post sample embedding. **a,** Detailed schematic overview of the generation of the histoarray from the histomold, followed by high-throughput embedding of CIVMs in the histoarray. **b,** Examples of three histoarrays filled with three different CIVMs pre-dehydration. **c,** Two histoarrays highlighted with different samples pre- and post-dehydration. For histomold abbreviations, see Suppl. Fig. 2.

**Suppl. Fig. 2:**
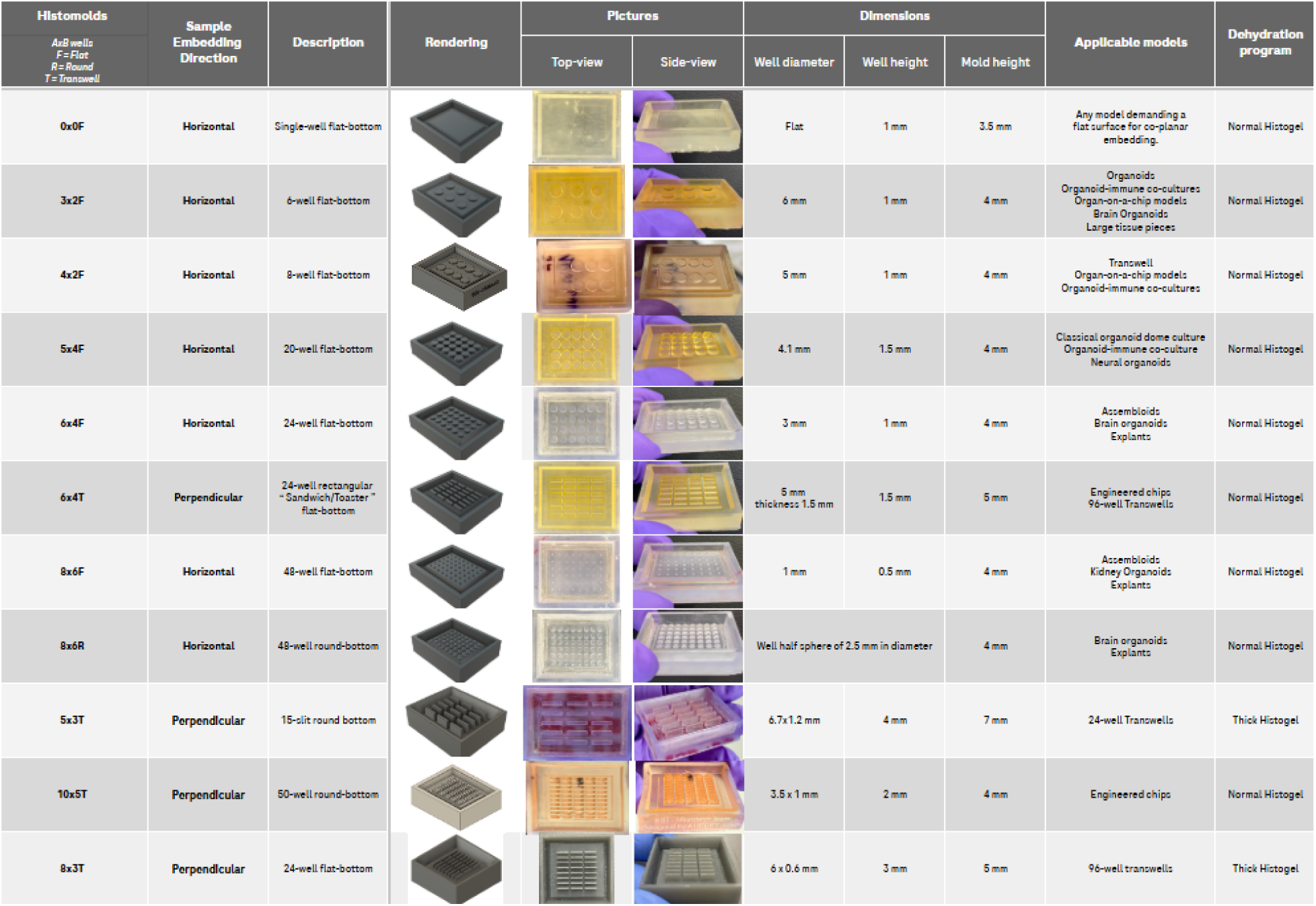
Design and application of current histomold landscape. Exhaustive list of in-house designed histomolds with various CIVMs and in-house use-cases per histomold.

By design, CIVMs can drastically differ from one another due to model-intrinsic and -extrinsic factors. These parameters directly steer the histomold design and histoarray-embedding workflow and, hence, demand different protocols per CIVM. To overcome this hurdle, we have designed a range of histomolds to facilitate simple and fast embedding of various CIVMs, while retaining a high density of wells per histomold (Suppl. Fig. 2). We grouped the CIVMs landscape into four workflows, aiming to develop overarching protocols governing the most common models and culture protocols in the field:

1. Single, mid to large-sized models (Fig. 1b-d)
2. Epithelial monolayers on transwells (Fig. 1e)
3. Soft ECM and suspension culture models (Fig. 1f)
4. Stiff ECM co-culture models (Fig. 1g)

The workflow is agnostic of the biological identity of the sample and almost exclusively determined by the respective intrinsic and extrinsic factors of the system (Table 1), and hence applicable to any CIVM. For example, assembloids (small), neural organoids (mid) or large tissue fragments (large) were embedded by simple sample transfer from the culture-well into the histoarray (Fig. 1b-d). To assess transwell-cultured epithelia in a perpendicular cross-section, detachment of the membrane can be easily performed, followed by simple transfer into the histomold (Fig. 1e). Organoids cultured within soft ECMs can be released into the supernatant prior to fixation, enabling them to be handled as suspension cultures, regardless of the organ of origin (Fig. 1f). For CIVMs that aim to understand the spatial dynamics between different cell populations within the microenvironment (e.g. epithelium-immune co-cultures), stiff ECM-embedded systems can be used, which enable embedding of the entire dome without perturbation of the spatial arrangement within (Fig. 1g). Although we cover a variety of CIVMs with a range of histomolds, the latter can be easily changed and extended to accommodate models not described within this manuscript utilizing our templates, such as organ-on-chip systems. Altogether, the high-content nature of the histoarrays spans various CIVMs, significantly reducing manual labor-intensive procedures while concurrently diminishing the consumption of expensive reagents. To illustrate these points, we showcase CIVMs of each aforementioned group and the respective workflows. We provide a designated figure per use case, sorted by ease of the respective embedding protocol, starting with single entities and concluding with immune-organoid co-cultures.

**Table 1:** Features of CIVMs that determine the design of the histomolds and the respective handling of the samples.

| Histomold Design Parameters | Examples |
| --- | --- |
| Size of Specimen | Small (<500 $\mu\text{m}^3$) vs. Mid (500-1000 $\mu\text{m}^3$) vs. Large (>1000 $\mu\text{m}^3$) |
| Entities per well | Single organoid vs. Multiple organoids |
| Culture system | Chip vs. (Plate-) Well vs. Transwell |
| Matrix | ECM vs. Suspension |

### High-throughput embedding of single, mid- to large-sized CIVMs

Cultures of animal or human-derived organs or tissue fragments *ex vivo* (explants) serve as models of the highest compositional complexity, preserving cell populations found *in vivo*, along with their spatial relationships, relative abundances and native extracellular matrix^49^. These models have recently been utilized to study immune cell activation in tumor tissues in response to specific immune checkpoint blockades^50^, chemotherapy^51,52^, targeted therapies^53,54^, exposure to oncolytic viruses^55^, assessment of fibrotic changes^56^ and studies of disease-specific mechanisms^57^. Whole-mount light microscopy enables high-throughput imaging assessment of samples; however, it remains challenging for large, cellular dense models, such as explants, due to insufficient light penetration. This limitation is typically remedied by FFPE-based imaging, albeit at the cost of throughput.

To address this, we designed a histomold with appropriate well dimensions typical of explants (8×6R histomold, Suppl. Fig. 2), enabling co-planar embedding of 48 samples within a single FFPE-block. To this end, we generated explants from patient-matched colon tumor and healthy colon tissue through manual scalpel tissue processing, followed by culture in 96-well plates to evaluate tissue integrity, morphology and composition at baseline and after 24 hours of explant culture, resulting in only two FFPE-blocks sufficient to cover a 96-well plate (Fig. 3a). After tissue preparation and 24 h culture, explants were fixed in 4% PFA in the culture plate, washed in PBS and individually transferred to a histoarray using tweezers (Fig. 3a). The use of tweezers minimized liquid carry-over; however, due to the spongy nature of tissues, additional liquid removal around the explants using a paper-wipe or air-drying was necessary before filling the histoarray with Histogel. Tissue morphology of the explants was assessed by H&E (Fig. 3b), which highlighted a systematic lack of organized crypt and villus structures in the tumor explant cohort, while normal explants showed a substantial presence of debris after 24 h in culture (Fig. 3b). mIF confirmed a wide heterogeneity in cellularity within the tumor explants, with certain fragments displaying packed and disorganized epithelial regions, while other fragments were composed predominantly of stroma, directly reflecting the organizational heterogeneity characteristic of tumor tissues (Fig. 3c). Both tumor and normal explants retained regions of epithelium, CD45^+^ cells (more abundant in normal than diseased), and CD31^+^ endothelial cells within the lamina propria, while overall tissue quality regressed over culture time (Fig. 3c).

**Fig. 3:**
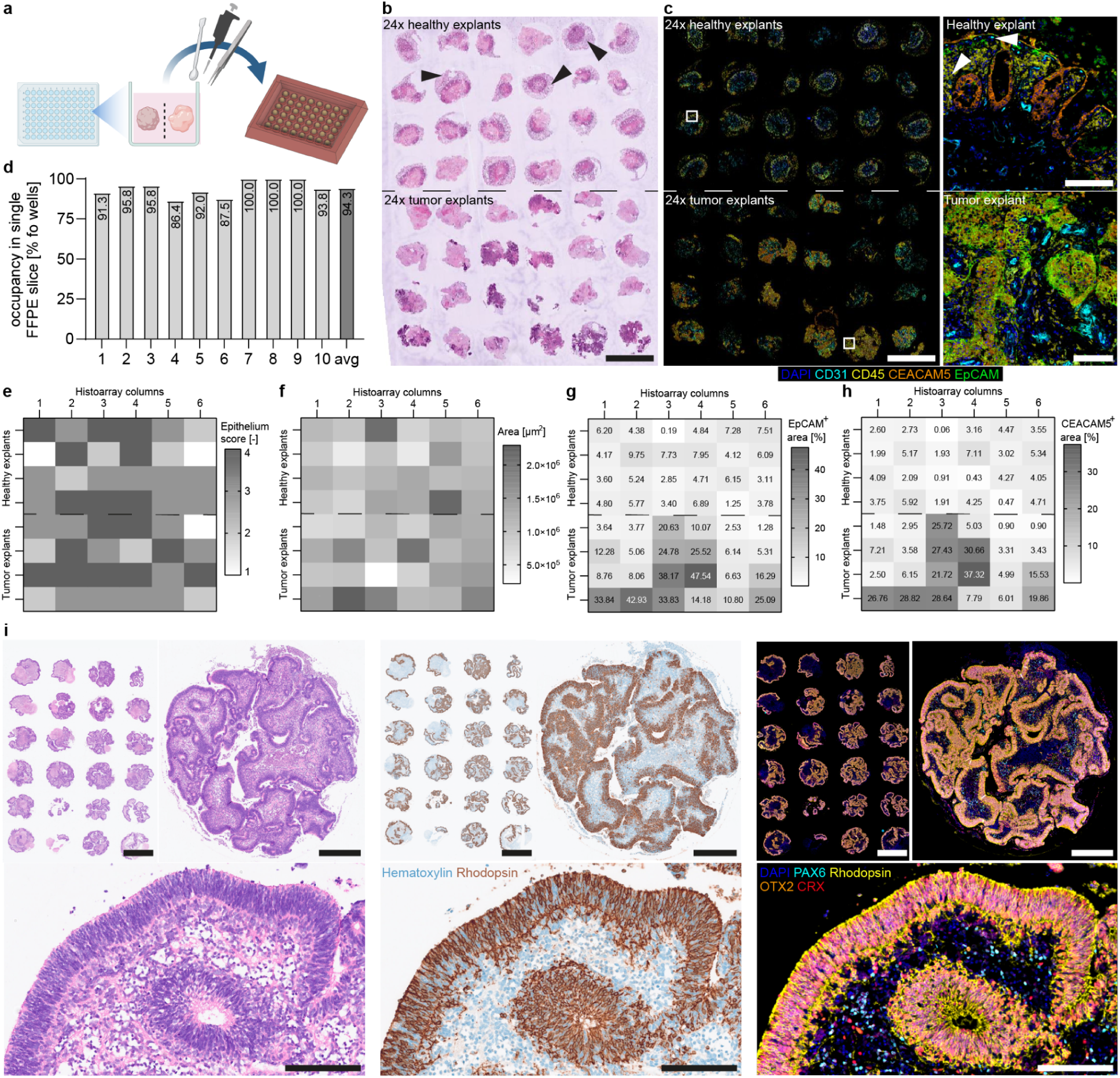
Histo-workflow and example for single, mid-large sized models: analysis of tissue fragments at high-throughput and differentiation assessment of murine retinal organoids. **a**, Schematic of the explant and retinal organoid embedding workflow by direct transferral of tissue fragments into an 8×6R histoarray. **b**, H&E image showing a full explant histoarray containing colon tissue after 24h ex vivo culture, comprising patient-matched colon normal explants (top half of the array) and colon tumor explants (bottom half). Black arrows indicate areas of non-cohesive tissue observed in normal explants upon culture (Scale bar: 2.5 mm). **c**, mIF image exemplifying normal and tumor tissue morphologies, respectively. White arrows indicate areas of debris (non-cohesive tissue areas) visible in normal explants upon culture (EpCam in green, CEACAM5 in orange, CD45 in yellow, CD31 in cyan and DAPI in blue). Histoarray overview scale bars: 2.5 mm. Magnified organoid scale bar: 100 µm. **d**, Explant histoarray well occupancy, quantified across 10 colon histoarrays, yielded an average of 94% of wells populated by explants in a given FFPE slice. **e**, Well-by-well scoring of epithelial content across 3 fresh colon explant TMAs. Epithelium content was found to be >50% of total well tissue content in 79.2%, 81.2% and 79.2% in TMAs #1,2 and 3, respectively (total average 79.9%). **f-h**, Quantification of explant cohesive tissue area (left) and EpCAM (center) and CEACAM5 (right) expression in matched normal and tumor explants upon 24h ex vivo culture, captured within one 48 well histoarray. EpCAM and CEACAM5 levels were normalized to the tissue area in each fragment. **i**, Consecutive sections stained by H&E, IHC and mIF of 28-day old in vitro generated murine retinal organoids in a twenty-four arrayed histomold (6×4F). Representative examples highlighted, with corresponding magnified inserts, demonstrating morphologically- and cell-type specific characteristics (photoreceptor cells-precursors (OTX2^+^, orange); photoreceptor nuclei (CRX^+^, red), rods (Rhodopsin^+^, yellow) and interneuron nuclei (PAX6^+^, cyan), including DAPI^+^ nuclei (blue). Histoarray overview scale bar: 2.5 mm. Single retinal organoid scale bar: 500 µm. Magnified retinal organoid scale bar: 100 µm.

As explants represent the full range of compositional heterogeneity of the tissue of origin, and further due to the manual preparation of fragments and potential residual mucus, explants show a considerable variability in sizes. This variability affected sample ‘glueing’, where these variations could result in suboptimal co-planar embedding. Regardless, the described workflow allowed us to achieve an average of 94% occupancy of histoarray positions by explants in sections (Fig. 3d). This occupancy metric improves when using tissues with homogeneous stiffness or lacking mucus (data not shown).

After isolating the mucosa from colon surgical resections, we assessed how robustly the epithelium was represented in the generated explants across three histoarrays (Fig. 3e; non-cultured). Specifically, the ratio of epithelium positive area to overall tissue fragment size was scored: a score of 1 indicated 0-25% epithelium positive area per fragment, 2 indicated 25-50%, 3 indicated 50-75% and 4 indicated explants with 75-100% epithelial coverage. We determined that the average probability of observing explants with more than 50% of the area positive for epithelium (i.e. a score of 3 or 4) was 79.9% (Fig. 3e). Together, these data indicate that the workflow yields explant histoarrays with nearly all wells occupied in any given FFPE section, and each well having a high probability of predominantly exhibiting epithelium.

This combination of metrics showcases the histoarrays as extremely powerful tools for streamlining the analysis of explants across experimental conditions. Additionally, sufficient sample replicates for image analysis of proteins of interest are easily generated: We utilized the dense sample-array section to quantify expression of targets for immunotherapy^58^, obtaining an overview of the variability in expression among multiple explants sourced from the same baseline tissue from a single patient, including matched normal and tumour tissue. Specifically, we quantified CEACAM5 and EpCAM expression through mIF, followed by quantification of antigen-expressing areas relative to respective cohesive fragment areas (Fig. 3f-h). Explants from colon tumor and normal tissue showed similar size distributions and, for this tissue donor, tumor explants displayed higher levels of both EpCAM and CEACAM5 compared to normal explants, with both groups showing high variability in antigen expression (Fig. 3g-h). Collectively, these data showcase the potential of the histoarrays around high-throughput embedding, staining and analysis of donor-matched healthy and diseased tissue explants, which is directly translatable to other large sized models.

Beyond the high-throughput advantages facilitated by histomolds, (histoarray-) FFPE blocks can be re-sectioned for future exploration without the need to re-grow specimens first. This is particularly valuable for models requiring lengthy maturation processes, such as neural or retinal organoids.

To illustrate this, we transferred a full twenty-four well plate of fixed mouse retinal organoids into the 6×4F histomold (Suppl. Fig. 2) to evaluate the morphology and differentiation status after 28 days of culture. Consecutive sections of the retinal organoid histoarray were subjected to H&E, IHC and mIF staining. At day 28, we found the neuroepithelium of the retinal organoids densely packed, polarized and stratified (Fig. 3i, H&E). Furthermore, morphologically-characteristic retinal layers including distinct retinal cell types were apparent by H&E. Compiling established single-IHC stains (representative of marker-establishment; Fig. 3i), performed on previous sections of the block, into a mIF panel we were able to confirm appearance of OTX2^+^ photoreceptor cell-precursors (PRCs) further differentiating into photoreceptor nuclei (CRX^+^), rods (Rhodopsin^+^) and interneuron nuclei (PAX6^+^) in distinct growth-media conditions (Fig. 3i). In conjunction with the explant example, the histomold facilitates easy co-planar embedding of these large CIVMs. Multiple consecutive sections can be generated and stained in parallel, increasing the sample coverage and robustness of the results significantly, at minimal processing, staining and imaging expense.

### Perpendicular cross-sections facilitated by histoarrays

Transwells are among the most common culture-vessels to study lung epithelial barrier integrity^59^, infection^60^, drug delivery^61^ and basic functions of barrier tissues^62^. Specifically, air-liquid interface (ALI) lung transwell systems are well established, as the ALI supports epithelial cell differentiation^63,64^. As these systems require multi-week differentiation protocols to differentiate into a mature epithelium, FFPE-based assessment is particularly valuable, as multiple sections can be obtained for both morphology and marker assessment. However, conventional perpendicular embedding of lung transwells is notoriously tedious, difficult and low-throughput, as it requires simultaneous precise placement of the transwell at 90°C in the metallic mold while pouring and polymerizing the paraffin in order to obtain a sufficient cross-section.

To overcome this, we designed histomolds with thin slit protrusions with the dimensions of standard 96-well and 24-well transwells that substantially ease the embedding of transwells (8×3T and 5×3T, Suppl. Fig. 2). Post fixation, we removed multiple sample-membrane constructs using either forceps to peel away the membrane (for 96-well transwells), a scalpel or biopsy puncher (for 24-well transwells) and transferred them directly to the histoarray, which guides perpendicular embedding by design (Fig. 4a). However, primary human alveolar epithelial cells cultures on 96-well transwells present particular challenges for FFPE processing, sectioning, and whole slide scanning, as the epithelium is only 1-2 cells thick and, thus, thinner than the underlying transwell membrane. Critically, we identified that the water bath temperature and incubation time on the water bath during sectioning significantly impacted sample integrity and section quality. Short incubation times of 5-10 seconds reliably yield high-quality H&E sections with straight samples and a majority of the epithelium in focus on scanned slides, regardless of water bath temperature (Fig. 4b). However, longer incubation times using a standard water bath temperature of 41°C result in frequent sample loss and progressive waviness in the sample-membrane constructs, with associated obscuration of the epithelium by the transwell membrane and intermittent regions that are out of focus. In contrast, using a lower water bath temperature of 36°C results in minimal drop off in section quality with up to 60 seconds incubation time. Thus, careful control of these variables is critical for enabling higher-throughput histologic evaluation of multiple transwells in one histoarray FFPE-block.

**Fig. 4:**
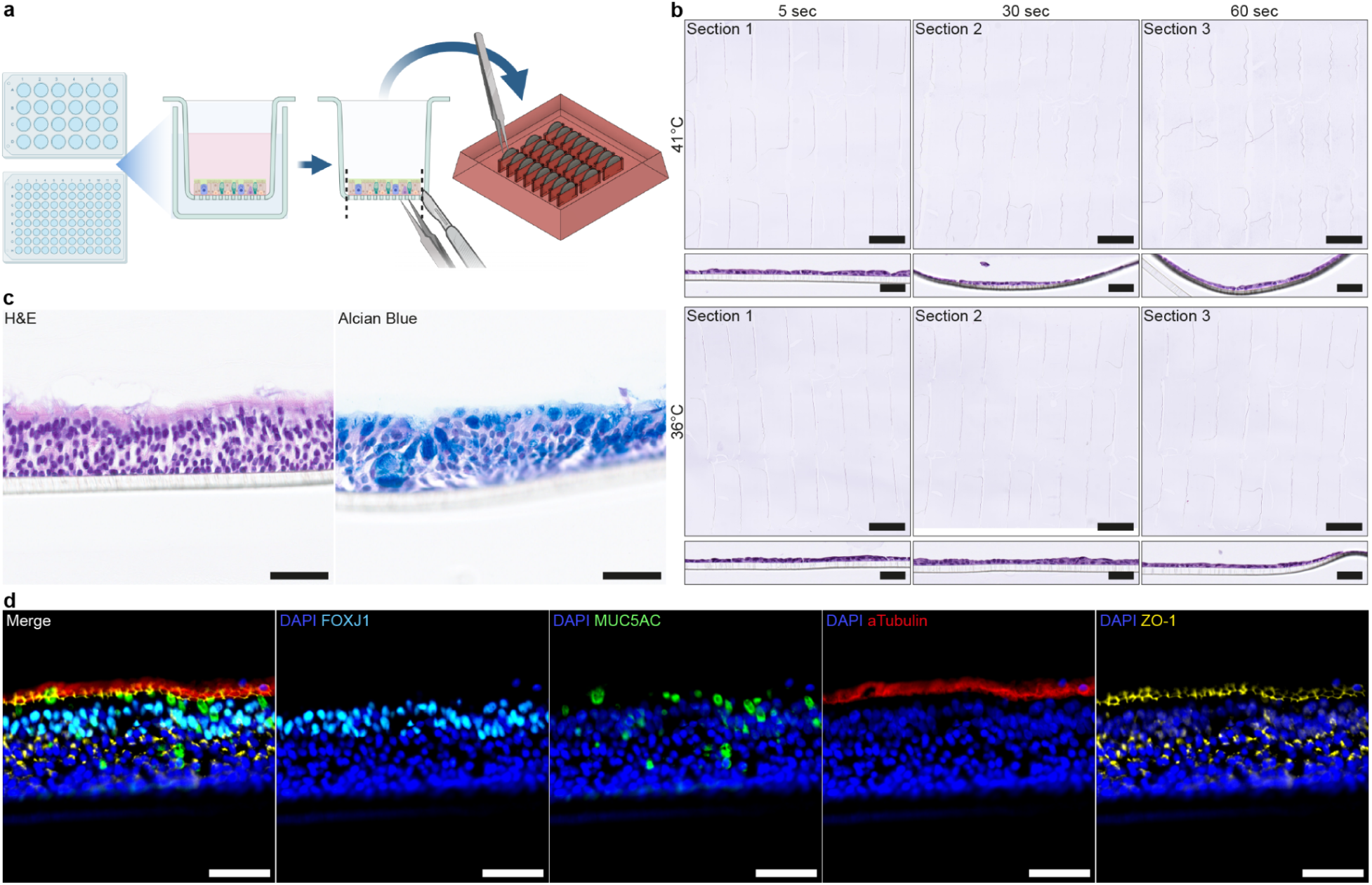
Histo-workflow and example for transwell-cultured in vitro models: perpendicular assessment of lung epithelium facilitated by the histomolds. **a,** Schematic overview of embedding lung transwells by peeling away the specimen-polymer complex with forceps (96-well transwells; 8×3T) or by removal with a scalpel (24-well transwells; 5×3T) and transfer into the histoarray. **b,** Consecutive sections of 10-day-old human primary alveolar epithelial cells cultured on transwells in an 8×3T histomold demonstrate the impact of water bath temperature and incubation time on the water bath on section quality. Representative magnified images of individual transwells illustrate that longer incubation periods at a standard temperature of 41°C lead to artifacts, including sample loss, sample folding, and irregular alignment, all of which impair the ability to capture all samples in focus during whole slide scanning (Scale bar: 2.5 mm; magnified insert scale bar: 40um). **c,** Representative H&E and Alcian blue images of a pseudostratified bronchial epithelium cultured on a transwell for 30 days (Scale bar: 50 µm). **d,** Representative mIF image highlighting the zonated, tight (ZO1^+^, yellow) and differentiated bronchial epithelium, exemplified by secretory cells (MUC5AC^+^, green) and FOXJ1^+^ ciliated cells (cyan) with ɑTubulin^+^ cilia (red) on the apical side of the epithelium, including DAPI^+^ nuclei (blue; Scale bar: 50 µm).

To assess a transwell model of bronchial airway epithelium, H&E and Alcian blue staining was performed, highlighting the pseudostratified epithelium rich in goblet cells (Fig. 4c). Using a consecutive section for mIF, we further confirmed the zonated architecture, exemplified by FOXJ1^+^ ciliated cells in the top layer of the epithelium, including ɑTubulin^+^ cilia on top of the polarized epithelium (Fig. 4d). Furthermore, strong apical ZO-1^+^ expression highlighted the tight barrier. Goblet cells (MUC5AC^+^) were mostly found in the upper layer of the epithelium, where they secrete mucus on the apical side of the tissue. These protocols demonstrate the utility of characterizing transwell models by routine pathology assays and could be easily applied to investigate lung epithelium from different regions of the airway (e.g. upper vs. lower), from diseased (e.g. COPD, cystic fibrosis) versus healthy donors, or from different treatment conditions, enabling easy and direct cross-comparison on the same slide. Furthermore, multiple sections at different depths of the histoarray could be utilized to strengthen the dataset, as the data per perpendicular section of an epithelial monolayer remains small. Although we focus here on the lung, these histomolds and workflows can be applied to other epithelial monolayers cultured on transwells, such as the intestinal epithelium.

### High-throughput embedding of soft ECM-embedded and suspension culture models

In the intestine, adult stem cells located at the crypt-bottom give rise to highly specialized, functional epithelial cell lineages, which is mirrored by intestinal organoids in vitro^65^. These three-dimensional, stem-cell derived organoids (adult and iPSC) are continuously maturing and differentiating in culture, recapitulating aspects of their parent organ given the right growth factors and time. In order to benchmark the status of epithelial differentiation of intestinal organoids from different intestinal regions and donors, assessment of cell-marker expression is critical to validate these models. However, as intestinal organoids are typically grown in a soft hydrogel (e.g. Matrigel), evaluation of such ECM-embedded systems via histology has proven difficult, primarily due to the challenging retrieval from the matrix. Particularly, upon in-situ fixation of the organoids in Matrigel, the matrix partially collapses, whereas some remains of the matrix stiffen and can easily adhere to the well or pipette tip. This may lead to the (partial) loss of the specimen and results in a low amount of biomaterial per section, not representative of the initial culture. Recently, protocols have been introduced that enable the culture of intestinal organoids in suspension^66^. Although protocols to embed such suspension cultures in FFPE-blocks are established, they remain a low-throughput workflow that does not facilitate co-planar embedding of multiple samples^38^.

Here, we demonstrate gentle liberation of soft ECM-cultured organoids by incubating the samples with Cell Recovery Solution for 40 min at 4°C, followed by harvest, fixation and direct dispensation into the 5×4F histoarray (Fig. 5a; Suppl. Fig. 2). Utilizing this approach, we were able to co-embed ten intestinal organoid donors in technical duplicates from various intestinal regions (Suppl. Fig. 5a). Using consecutive sections for H&E, IHC and mIF, we were able to assess morphological integrity by H&E, expression of a cancer immunotherapy-relevant target protein^67–69^ (IHC) and heterogeneous organoid differentiation (mIF) from a single FFPE-block (Fig. 5b; Suppl. Fig. 5c).

**Fig. 5:**
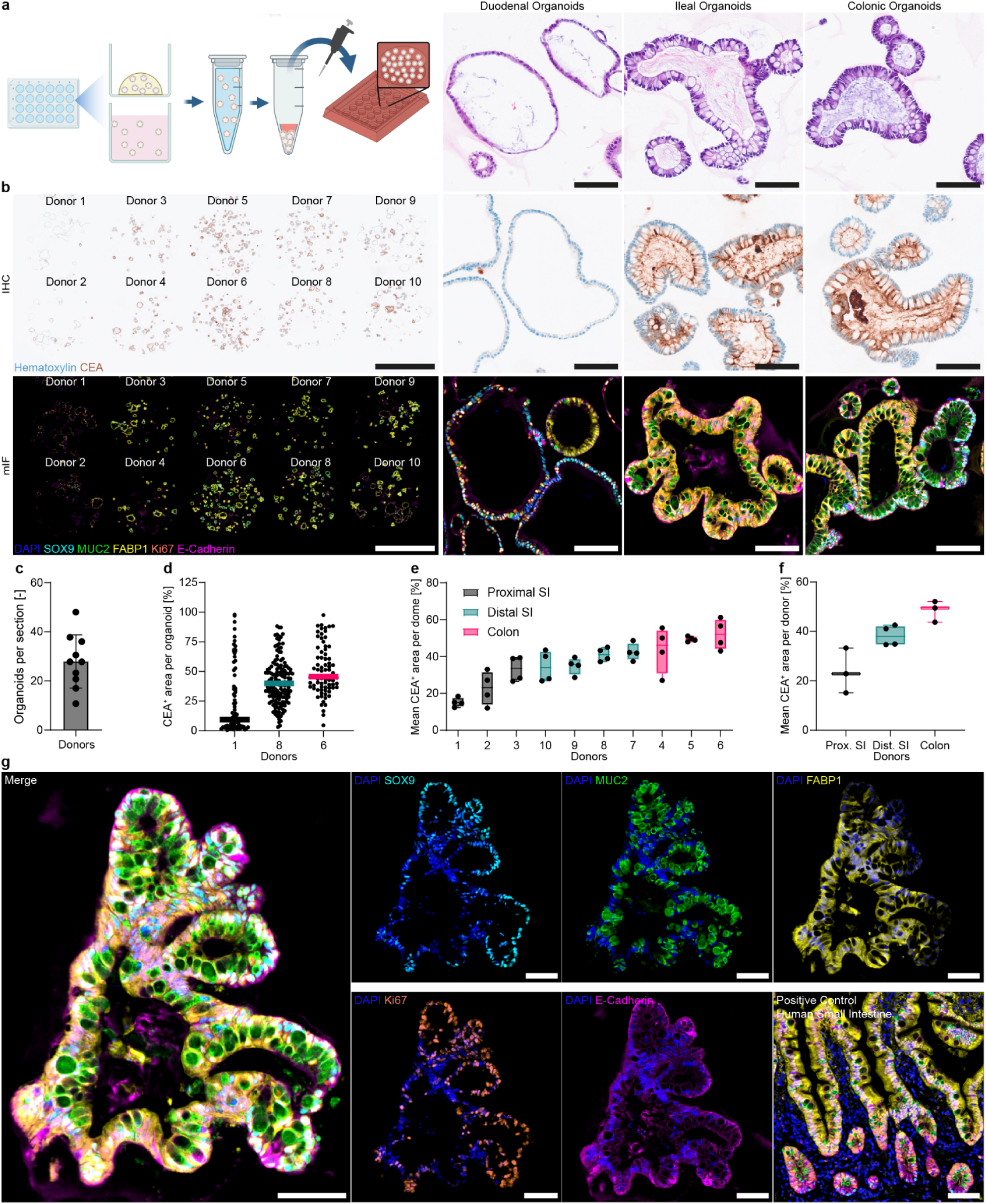
Histo-workflow and example for soft-ECM embedded and suspension in vitro models: differentiation assessment and cancer immunotherapy target expression of ten organoid donors across the GI-tract. **a,** Schematic overview of processing soft ECM-embedded organoids (e.g. Matrigel) by organoid recovery from matrix and/or suspension-cultured organoids, followed by concentration of samples and dispensation into a histoarray. **b,** Representative H&E, IHC and mIF images of half of the 5×4F histoarray across all donors, including dozens of 6-day old organoids per well (Scale bar: 2.5 mm). Corresponding magnified inserts highlight representative organoids from different regions, demonstrating a polarized (indicated by strong apical CEA^+^ expression (brown)), diverse and mature epithelium (stem-cell associated cells (SOX9^+^, cyan), goblet cells (MUC2^+^, green), enterocytes (FABP1^+^, yellow), proliferating cells (Ki67^+^, orange), adherens junctions (E-Cadherin^+^, pink), DAPI^+^ nuclei (blue). Magnified inserts scale bar: 100 µm. **c,** Mean of organoid number per section across the ten donors. **d,** Quantification of CEA^+^ area of all organoids in one histoarray-well of three representative donors. **e,** Quantification of the mean CEA^+^ expression of the organoids per well across two sections per donor. **f,** Mean CEA^+^ expression across two sections per donor respective of the region of origin. **g,** Example of a 6-day old ileal intestinal organoid, showcasing differential marker expression between a crypt-like area (Ki67^+^, SOX9^+^, FABP1^-^) and a villus-like structure (Ki67^-^, SOX9^-^, FABP1^+^), including duodenal tissue as positive control (Scale bar: 100 µm).

**Suppl. Fig. 5:**
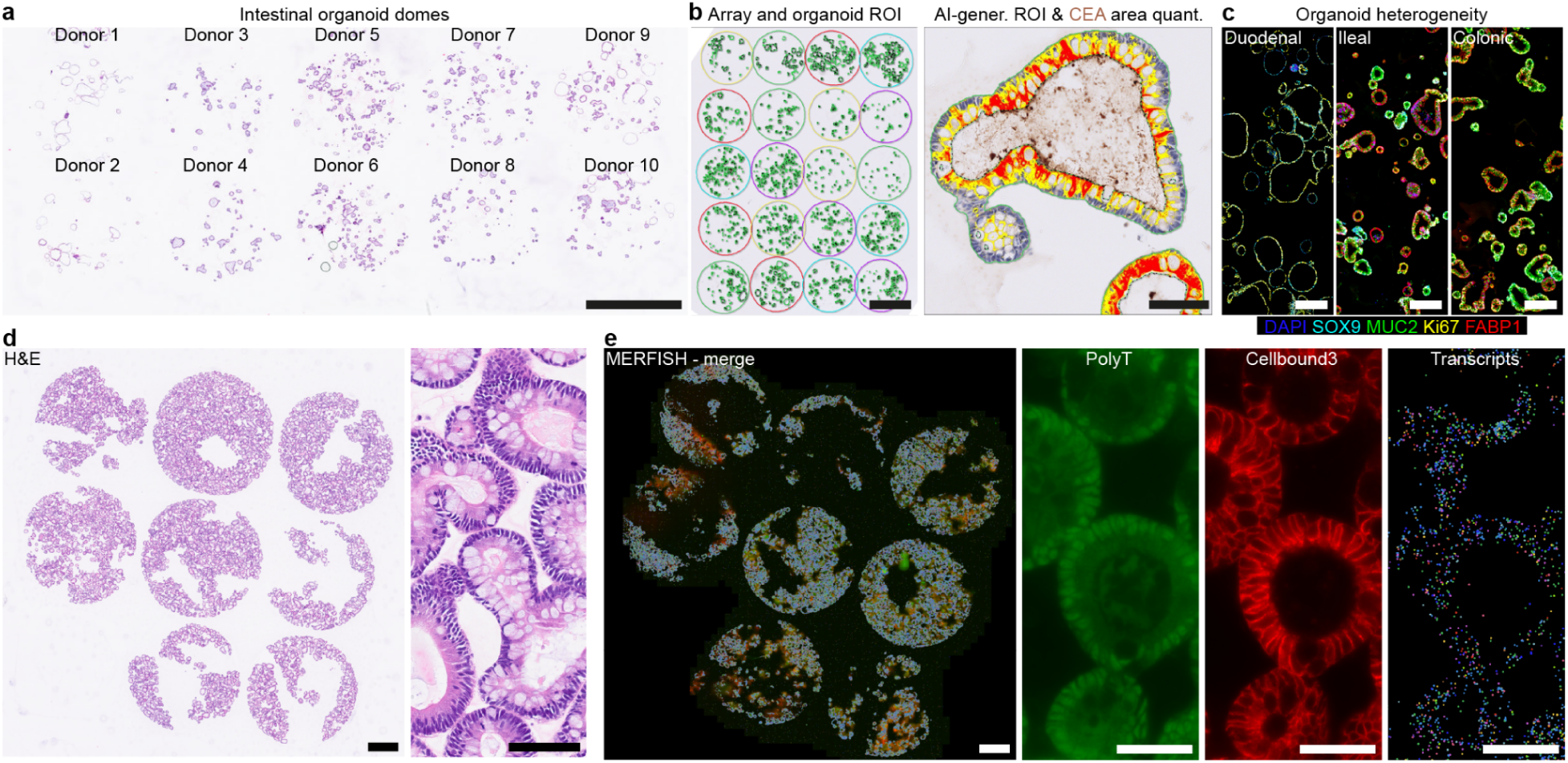
Analysis of single organoids in histoarray and proof of concept of MERFISH on samples in histoarray. **a,** Representative H&E of half of the 5×4F histoarray, showcasing dozens of 6-day old organoids per well (Scale bar: 2.5 mm). **b,** Annotation of each well as an individual sample (Scale bar: 2.5 mm). Utilization of a RandomForest classifier (HALO AI) to automatically detect and annotate each organoid as a single object, separating lumen and epithelium. Quantification of the CEA^+^ ROI within the epithelium of the organoid (Scale bar: 50 µm). **c,** Representative mIF image exemplifying the heterogeneous intra-region/donor and inter-region/donor intestinal marker expression on one slide (Scale bar: 250 μm). **d,** H&E staining of intestinal organoids embedded in 8 wells of a flat-bottom 5×4F histoarray (Scale bar: 1 mm). **e,** Consecutive slide from c) subjected to MERFISH spatial transcriptomics (Scale bar: 1 mm). Overview and representative magnified images of PolyT signal (total mRNA, GREEN), cell boundary staining (red) and transcripts detected (colorful dots) with a custom panel of gene probes against duodenal intestine cell markers (Scale bar: 50 μm).

Given the arrayed layout of the samples, virtual TMA-annotations were swiftly created to quantify CEA expression per organoid of each donor, utilizing a RandomForest algorithm (Suppl. Fig. 5b). Across the ten different donors, a mean of 28 organoids per section was detected (Fig. 5c). Despite this number being sufficient for analysis, the density of the organoids per well can be further increased to enhance biological content per section (Suppl. Fig. 5d). CEA expression was visualized for all identified organoids of three representative donors (Fig. 5d), highlighting the distribution of expression within a technical replica. We then merged the mean of all technical replicas per donor to find that colon organoids have the highest CEA expression level (Fig. 5e), reflected by region specific visualization (Fig. 5f). Next, we assessed the heterogeneous differentiation status of the organoids by applying 6plex mIF (Suppl. Fig. 5c), highlighting the potential of histomolds to drastically reduce the expense of reagents when subjected to costly stainings. While ileal and colonic organoids differentiated well within the culture period (Fig. 5b), highlighted by ubiquitous FABP1^+^ enterocytes, MUC2^+^ goblet cells and thick, polarized epithelium (HE and E-Cadherin), such degree of differentiation was not observed in the duodenal organoids (Fig. 5b); neither via morphological features nor on protein level. As previously reported, intestinal organoids mimic the spatial patterning of their organ of origin in vitro^70^. We were able to confirm this by direct comparison to primary duodenum tissue (Fig. 5g), including crypt-like buds (FABP1^-^, Ki67^+^, SOX9^+^) and a ‘villus-like’ area (FABP1^+hi^, Ki67^-^, SOX9^-^).

In addition, we assessed whether our histo-workflow is amenable to sensitive spatial transcriptomics methods. To this end, we subjected a small but dense histoarray (8 wells) with intestinal organoids to multiplexed error-robust fluorescence in situ hybridization (MERFISH) for FFPE samples (Suppl. Fig. 5e). We confirmed that mRNA quality after histoarray processing was suitable for spatial transcriptomics (DV_200_ 71%, data not shown). In addition, we were able to detect hundreds of transcripts outlining the epithelium of the intestinal organoids (Suppl. Fig. 5e), albeit transcripts per cell being low, demanding further optimization in the future.

Nonetheless, this feasibility study for spatial transcriptomics widens the application of the histomolds vastly. Crucially, the histomolds would, by design, alleviate the typically tedious placement and section-arrangement within a small imageable area of these extremely costly spatial omics techniques.

### Stiff ECM-embedded co-cultures to retain spatial information

With the rise of immunotherapy for a range of immune-mediated pathophysiologies^71–73^, the concomitant contribution of the tumor microenvironment (TME) and spatial context has become increasingly more evident^74–76^. Although adult stem-cell derived organoids represent a great tool for disease modeling, they are devoid of critical tissue compartments such as the immune system, which is fundamental for modeling and optimizing the effects of immunotherapy. Recently, multiple groups have started to co-culture immune cells with organoids to overcome this gap^77–81^. However, these cultures are typically subjected to destructive methods such as flow cytometry or the soft-ECM (e.g. Matrigel) disintegrates upon fixation, thereby losing the spatial context and information about the TME^82^. Although live imaging enables in situ assessment of the system, it is limited by the amount of markers and demands complex analysis to utilize the large data^77^.

To overcome this gap, we’ve recently engineered a stiff collagen-Matrigel matrix that withstands fixation and, hence, enables the assessment of immune-organoid interactions in the context of the microenvironment^83^ (Fig. 6a). Briefly, after fixation of the sample in the well, Histogel™ is dispensed and polymerized on top of the dome inside the plate. Afterwards, the dome-Histogel™ construct can be retrieved, excess Histogel™ removed and the sample transferred to a 5×4F histoarray (Suppl. Fig. 2). By utilizing this protocol to retain spatial integrity of co-cultures with the histomolds, we aimed to assess the spatiotemporal anti-tumor effect of an EpCAM-targeting T cell bispecific (TCB) on PBMC-tumoroid co-cultures within one FFPE-block.

**Fig. 6:**
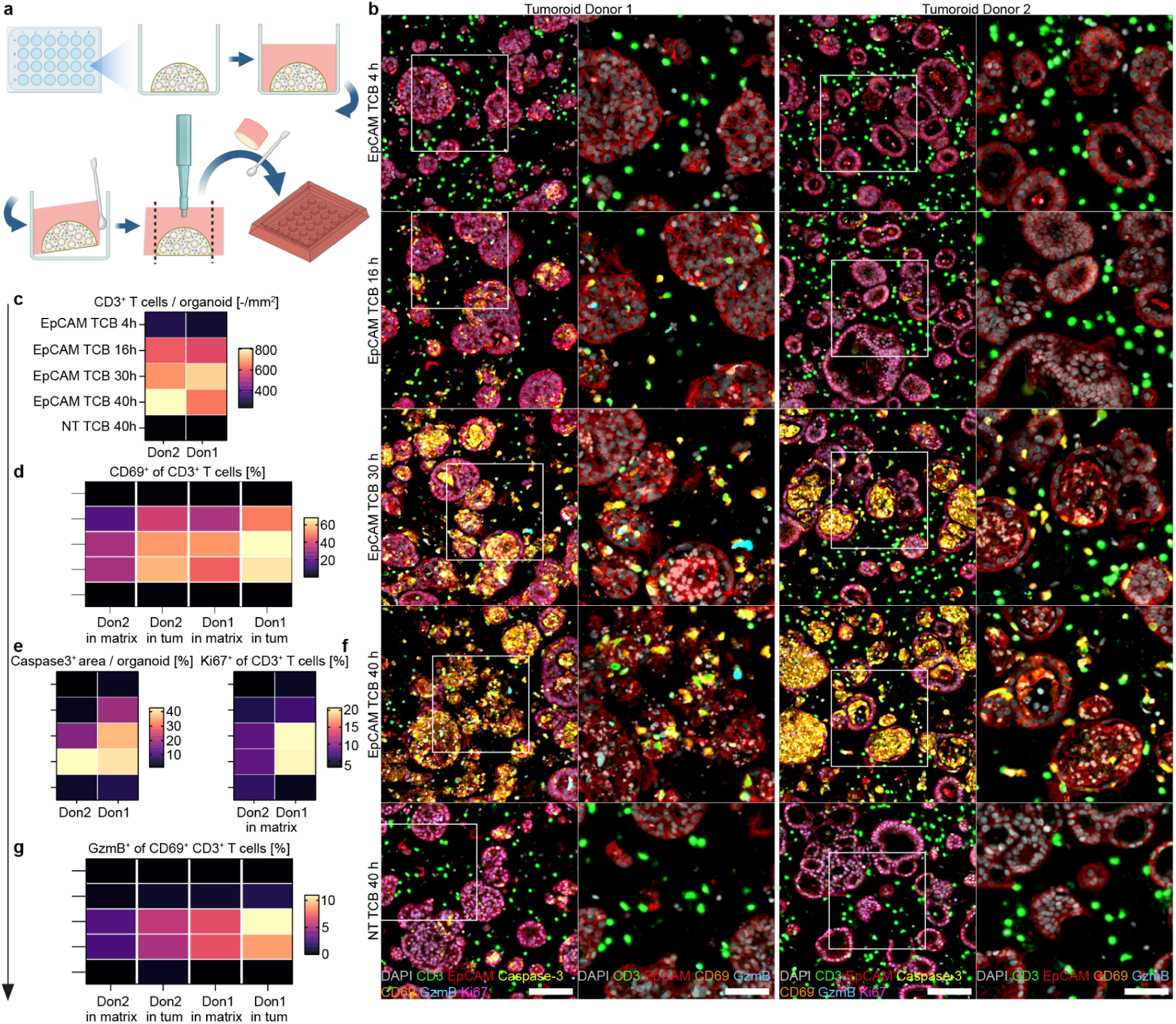
Histo-workflow and example for stiff-ECM embedded in vitro models: spatiotemporal analysis of T cell bispecific efficacy assessment on immune-tumoroid co-cultures. **a,** Schematic overview of embedding stiff ECM-cultured (e.g. Collagen-Matrigel) PBMC-intestinal tumoroid co-cultures to retain spatial organization by in-well Histogel™-polymerization, followed by lifting and punching out the co-culture dome and transfer to the 5×4F histoarray. **b,** Representative 7plex mIF images following the activation of T cells and subsequent cytolysis of tumoroids post EpCAM TCB treatment (5 µg/mL) per donor and time points from the same histoarray section (Scale bar: 100 µm). Magnified inserts highlight T cell infiltration and activation within cancerous epithelium (Scale bar: 50 µm). **c-g,** Quantification of EpCAM TCB-triggered CD3^+^ T cell infiltration (green) per tumoroid (EpCAM^+^, red), followed by upregulation of activation marker CD69^+^ (orange) and cytotoxic granules (GzmB^+^, cyan) resulting in caspase-3^+^ apoptosis (yellow) of the tumor epithelium, induction of CD3^+^ T cell proliferation (pink), including DAPI^+^ nuclei (grey).

**Suppl. Fig. 6:**
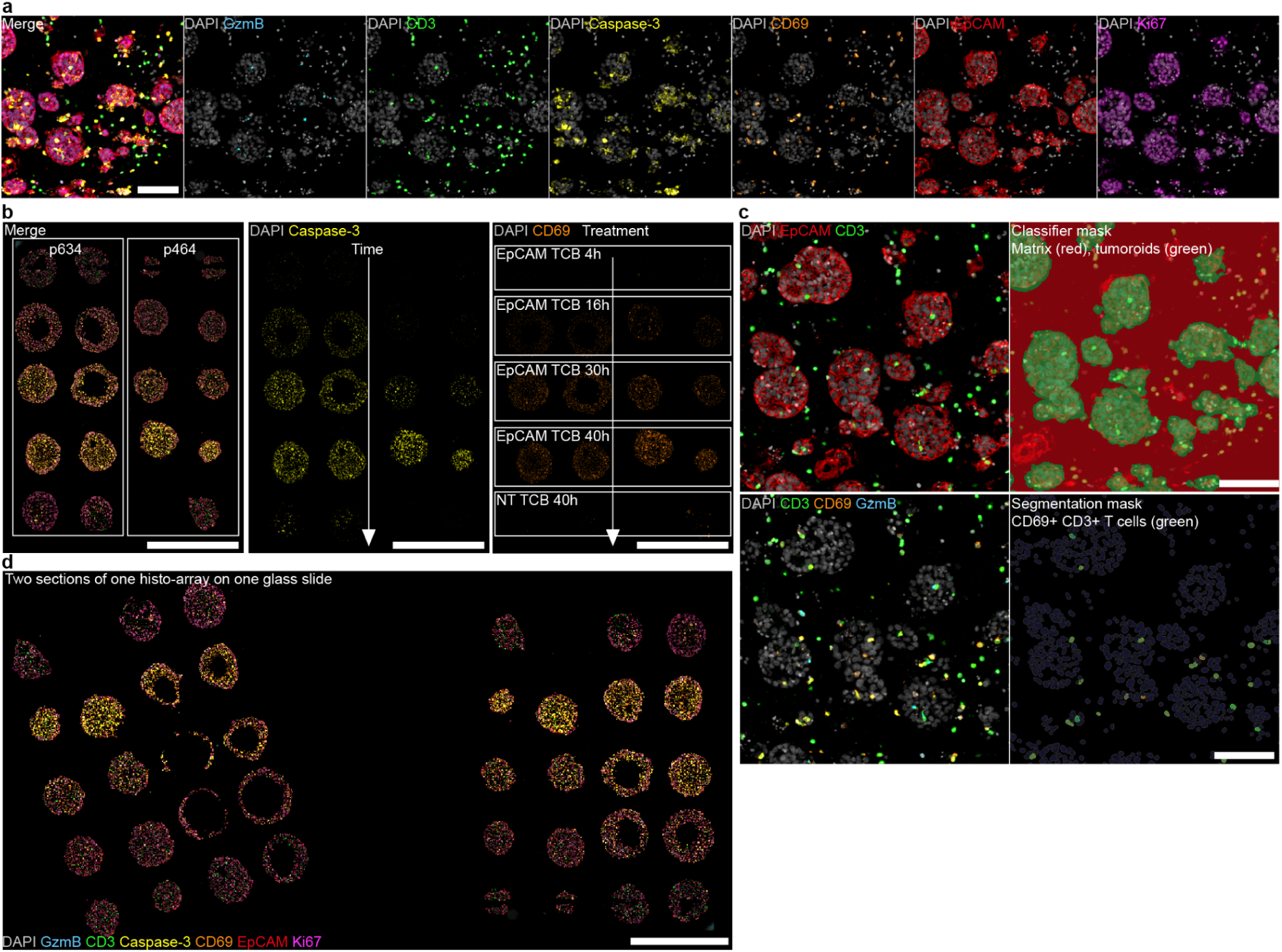
Analysis of T cell bispecific time course of two immuno-tumoroid donors on two sections at different depths on one glass slide. **a,** Representative 7plex mIF image, exemplifying each channel separately. Scale bar: 100 µm. **b,** histoarray overviews indicate EpCAM TCB-mediated T cell activation (CD69^+^, orange) and killing pattern (Caspase-3^+^, yellow) across donors and time (Scale bar: 5 mm). **c,** Utilization of a RandomForest classifier (HALO AI) to automatically detect and annotate each tumoroid as a single object (green mask). Analysis and segmentation of the CD69^+^ CD3^+^ T cells in both tumoroids and matrix (Scale bar: 100 µm). **d,** Two sections of the same histoarray at different depths to increase technical sampling size and statistical power on the same slide. Asymmetric sample embedding enables swift orientation of section and sample identification (Scale bar: 5 mm).

To this end, we treated the co-cultures (two tumoroid donors) with 5 µg/mL of EpCAM-TCB across four time points (in duplicates) and transferred the resulting 20 stiff-ECM domes into a single 5×4F histoarray (Suppl. Fig. 2). By utilizing the co-planarity of the samples in the histoarray, we generated two sections at different depths and transferred them onto one glass slide to increase robustness of the resulting data (Suppl. Fig. 6d). Deploying a 7-plex mIF approach (Suppl. Fig. 6a), we first assessed the cultures at low magnification to get a quick overview of the outcome of the experiment. Thereby, we were able to observe increasing apoptosis induction (Caspase-3^+^ cells) in the tumoroids (EpCAM^+^) and strong co-expression of CD69 (activation) in the CD3^+^ T cells in a temporal manner across both donors (Suppl. Fig. 6b). Next, we used a RandomForest algorithm to classify tumoroids and the microenvironment as separate regions of interest (ROI) and determined the T cell phenotype within both (Suppl. Fig. 6c).

We found that the T cells increasingly infiltrate the tumoroids (Fig. 6b), become activated (CD69^+^ CD3^+^), upregulate cytotoxic GzmB^+^ granules and start to proliferate (Ki67^+^), leading to the induction of apoptosis (Caspase-3^+^) of the EpCAM^+^ tumoroids (Fig. 6b-g). Interestingly, the T cells within the tumoroids exhibited higher levels of CD69 and GzmB than the ones remaining in the matrix, suggesting TCB-triggered, local response by the T cells that are in close proximity to the antigen followed by the activation of the T cells in the periphery (matrix; Fig. 6b, d, g). This highlights the critical need for evaluating such co-cultures in the context of cancer immunotherapy by spatial methods. Beyond the potential of the histoarray to consolidate an entire TCB-efficacy assessment longitudinally, the histoarray enabled simultaneous comparison of the effects on two tumoroid donors. Unlike destructive methods, the FFPE-block of this experiment could be re-sectioned and subjected to more stainings, potentially elucidating novel spatial mechanisms of action or deeper immune-phenotyping. Using this approach, we were recently able to capture differential infiltration and killing patterns of the T cells in the context of healthy and cancerous epithelium from the same donor^83^. Furthermore, we’ve utilized this approach to study the intraepithelial character of tissue-resident memory T cells when co-cultured with autologous organoids at homeostasis^84^, opening new avenues to study epithelium-immune interactions *in vitro*.

## Discussion

The field of CIVM research is rapidly evolving and opens new avenues for basic and translational research in tissue development, function and disease. With the legal removal of animal testing as a necessity to move molecules forward in pharmaceutical development (FDA Modernization Act 2.0^35^), it has become paramount to incorporate CIVMs at each stage of preclinical development. Therefore, in-depth and systematic characterization of these models is pivotal. As histological data has been part of the standard repertoire for regulatory authorities for decades, it is logical to apply these protein-based readouts to CIVMs to explore their spatial biology and support regulatory decision making. However, in order to facilitate the large sample volumes of the CIVM research field, protocols have to be developed to overcome the traditionally very low-throughput and highly manual-labor intensive histopathology workflows. Although some previous work on arrays for CIVMs has been published, the focus has mostly been on large, spherical tissues, which are comparably simple to handle^45–48^.

Here, we developed a holistic histo-workflow, that overcomes multiple shortcomings of traditional histo-processing and showcases the great potential of histopathology for CIVMs. In particular, we’ve generated a large battery of histomolds designed to enable high-throughput embedding of a wide range of CIVMs of various sizes, morphologies, culture-vessels and methods (Table 1). Our 3D-printed histomolds can be easily designed, manufactured and adapted, while requiring relatively low expertise and inexpensive equipment. The transfer of samples is swift, once routinely applied, enabling co-planar embedding of up to 48 samples in one FFPE-block. Determined by the depth of the sample in the histoarray, dozens (e.g. dome co-cultures) to hundreds of consecutive sections (e.g. transwells) can be obtained and stained to interrogate the models. To highlight the throughput and range of readouts of the workflow, we subjected consecutive sections to different staining techniques using automated stainers and obtained rich datasets of our CIVMs, while significantly reducing cost of reagents. Without the use of automated stainers, the benefit of using histomolds is retained and potentially more impactful, as it substantially decreases manual processing time. Furthermore, the arrayed layout eases image analysis, as intra- and inter-staining variability is minimized, ROIs can be instantly generated and analysis can be run much faster, due to numerous samples being present within the same image. Additionally, various types of negative controls can be co-embedded by experimental design, further simplifying detection scoring or identification of unspecific antibody binding. Moreover, we demonstrated that the dense sample-array facilitated by the histomolds is ideal to accommodate highly advanced and costly spatial transcriptomics methods such as MERFISH, broadening the applicability of the method. Lastly, as FFPE-samples can be stored for years, the specimen can be re-sectioned and re-stained whenever needed, without the need of (re-)generating the CIVMs.

### Limitations of the study

Although the histomolds offer great promise and are generally easy to use and implement, the process of embedding and subsequent sectioning can be challenging, particularly in the case of small, spherical models (<0.5 mm^3^) that are intended to be embedded in individual wells (e.g. 8×6R; Suppl. Fig. 2). Consequently, small irregularities such as ‘non-glueing’ them into the well prior to filling up the histoarray, transfer of wet samples into the histoarray, slight misalignment of the microtome cutting angle or dehydration artifacts due to histoarray overfilling (resulting in the absence of a flat surface) might negatively affect the total coverage of embedded samples in the sections. To avoid this, pooling the specimen by resuspending multiple technical replicas in Histogel and directly dispensing them into e.g. the 6×4F histoarray (Suppl. Fig. 2) will generate deeper sectioning volume and more biological material per well. Regardless of the CIVM, the microtome has to be perfectly calibrated at a 0° angle. Dehydration errors can be easily avoided by sufficient pre-dehydration in formalin as well as by minimizing the thickness of the histomolds in the design phase, as we found larger volumes of Histogel to correlate with dehydration issues (extreme shrinkage or stiff arrays post-dehydration). Since CIVMs are more challenging to section and analyse compared to more traditional and larger organs, it is critical to train and inform histopathology staff about the specific samples (e.g. size of the CIVM), special sample handling needs, the layout and how many sections to generate. Lastly, although the process to generate our histomolds is significantly easier than previous workflows^43,45,46^, it still requires some basic 3D printing knowledge. However, with the templates provided in this manuscript (Suppl. Fig. 2), the histomolds can be directly printed (or easily adjusted) and integrated into any histology-lab routine.

Despite these potential limitations, the data provided here illustrate how histo-workflows can be applied to CIVMs by integrating them into the standard histology pipeline through the use of histomolds, yielding highly informative datasets in a standardized fashion. We hope that the described workflows encourage CIVM researchers to consider histopathology as a high-throughput readout for high-quality imaging data acquisition.

## Methods

### Ethics statement

Human tissues and associated clinical information, along with concurrent data collection and experimental procedures, were obtained from patients undergoing tumor resections within the framework of the non-profit foundation HTCR (Munich, Germany) and at the University Center for Gastrointestinal and Liver Disease (Clarunis; Basel, Switzerland) as well as Humanitas Research Hospital (Milan, Italy), including informed patient consent. The HTCR Foundation’s framework received approval from the ethics commission of the Faculty of Medicine in the Ludwig Maximilian University (no. 025-12) and the Bavarian State Medical Association (no. 11142). The framework of the University Center for Gastrointestinal and Liver Disease was approved in accordance with the Helsinki Declaration and reviewed and approved by the ethics committee (Ethics Committee of Basel, EKBB, no. 2019-02118). The framework of the Humanitas Research Hospital received approval from the ethics commission Comitato Etico Terrirotiale Lombardia 5 (Ethical approval 3631).

### Complex in vitro model culture

#### Explants

Patient-derived surgical tissue resections were received the morning after surgery. In order to preserve tissue viability, all tissue processing was carried out on ice with addition of sufficient wash medium (advanced DMEM/F12 + 1:250 Primocin + 2% PenStrep), to prevent drying out of tissue during manipulation. As a first step, the abundant mucus layer atop the epithelium was gently scraped off using the dull side of a scalpel blade. Secondly, because a full-thickness surgical resection of colon includes mucosal, submucosal and inner and outer muscular layers, the mucosa and submucosa were resected from the remaining tissue prior to fragmentation into explants, by careful separation using a scalpel. Once the relevant layers were selected, explants were generated by cutting the tissue further in the x and y directions until fragments of 1 mm^3 size were obtained. Explants were stored in wash medium on ice until the desired number of fragments were cut, at which point they were distributed into flat-bottom 96 well plates for 24 hr ex vivo culture in culture medium (advanced DMEM/F12 + 10% FBS + 1X Glutamax + 1:250 Primocin + 2% PenStrep) at 5% CO2 and 37 degree C. At culture endpoint, explants were fixed in 4% PFA directly in the culture plate (see ‘Fixation & Harvest Method III’), washed in PBS and transferred to a de-molded histoarray using tweezers in a one-by-one fashion. Transfer with tweezers minimized carry-over of liquid; however, due to the spongy nature of tissues, additional removal of liquid around explants using a kim wipe or air-drying was necessary, prior to filling the histoarray with histogel (see ‘Transfer Method I’).

#### Neural Organoids

Mouse neural organoids were derived following an adapted version of Lancaster et al. (Nature Protocols 2014)^85^ using the murine ESC line SBR.

#### Transwells

Refer to Swart et al. (Nature Microbiology)^60^ for the detailed protocol of the normal human bronchial transwell culture (Catalogue #CC-2540, Lot#18TL346815, Lonza).

Alveolar transwell cultures were established using a protocol adapted from an alveolus lung-chip protocol^86,87^. Briefly, human primary alveolar epithelial cells (HPAEC; Cell Biologics, Cat# H-6053, Lot# F101517Y72) from normal lung tissue were expanded to passage 6 (P6) using SAGM™ Small Airway Epithelial Cell Growth Medium (Lonza, Cat# CC-3118) per the manufacturer’s instructions with modifications. Three days prior to seeding, HPAECs were thawed and expanded to P8 in SAGM complete medium, which consists of SAGM supplemented with 5% heat-inactivated fetal bovine serum (HI-FBS; Sigma, Cat# F4135). HTS 96-well transwells (Corning, Cat# 7369) were coated for 30 minutes at room temperature with an extracellular matrix mixture containing 0.03 mg/mL fibronectin (Corning, Cat 356008), 0.2 mg/mL collagen type IV (Sigma, Cat# C5533) and 0.005 mg/mL laminin (Sigma, Cat# L6274) in endotoxin-free calcium/magnesium-free phosphate buffered saline (PBS; EMD Millipore, Cat# TMS-012-A), then equilibrated with SAGM complete medium in the basolateral compartment. HPAECs were seeded at 1.25 x 10^5^ cells/mL (100 µL) on the apical surface and incubated overnight at 37°C in a 5%, humidified CO_2_ incubator. The next day, media in both compartments were replaced with SAGM complete media supplemented with 100 nM dexamethasone, 5 ng/mL keratinocyte growth factor (KGF; ThermoFisher Scientific, Cat# PHG0094), 50 μM cyclic adenosine monophosphate (cAMP; Sigma, Cat# B7880) and 25 μM isobutyl methylxanthine (IBMX; Sigma, Cat# I7018). On Day 3, the apical medium was removed to establish an air-liquid interface and the basolateral medium was switched to DMEM/F-12 (Gibco, Cat# 11320033) with 1% pencillin-streptomycin (Gibco, Cat# 15140122), 1X GlutaMAX™ (Gibco, Cat# 35050061) and 10% HI-FBS. Medium was refreshed every 3-4 days. At Day 7 post establishment of an air-liquid interface (10 days in culture), cells were washed with PBS and fixed in 4% paraformaldehyde (PFA) for 1 hour at room temperature before storage in 70% ethanol for FFPE processing.

#### Intestinal organoids

Refer to Fuji et al. (Cell Stem Cell 2018)^70^ for the detailed protocol of the human intestinal organoid culture.

#### Intestinal tumoroid co-culture

Refer to Harter at al. (NBME 2024)^83^ for the detailed protocol of the human intestinal tumoroid, immune and tumoroid-immune co-culture.

#### Retinal mouse organoids

Refer to Völkner et al. (Stem Cell Reports)^88^ for the detailed protocol of the mouse iPSC-derived retina organoid protocol.

### Histomold design and manufacturing

The mold geometry comprises an array of cylindrical wells with a flat or round bottom sharing the same plane of reference. Surrounding this array, a wall made of the same material (cast gel) allows the pouring of gel over the organoids once they are in place. Filling the mold with gel until the wall height results in a uniform block of gel with a top flat surface.

To obtain this design, the casting mold contains the following features (see profile):

- A thick base of lateral dimensions (A-B) 37-31 mm and a minimum height of (I) 12 mm to obtain enough rigidity to ensure a flat 3D printed mold
- A surrounding wall that will contain the first gel to be subsequently demolded. This wall should have a height of (H) 4 mm,and a thickness of (C) 2 mm at its top. An angle (F) 100 °facilitates the demolding of the gel.
- A groove with respect to the middle surface that will create the wall of the demolded gel. This groove has a depth of (J) 2 mm and is (M) 2 mm wide at the base. As before, an angle of (G) 60 ° facilitates its demolding.
- A middle flat surface with lateral dimensions (D-E) 27-21 mm and protruding cylinders, cuboids or half disks that will serve to form the wells. The exact well sizes used are described in Suppl. Fig. 2 every specific design. In general, the well diameter ranges from 0.25 to 5 mm, and the height from (K) 0.25 to 1.5 mm. For cylinders, the top can be flat or round. In the latter case the round part is a semisphere of radius equal to cylinder radius. The distance between the top of the cylinder/cuboid/disk and the external wall top should be at least (L) 1 mm, to ensure the integrity of the final molded part.

The three dimensional design of the mold was defined by computer aided design software (Autodesk Fusion 360). We additionally provide in supplementary the 3D file of the designs described in this work. The design was imported into a 3D printer-compatible software, in our case, PreForm from Formlabs, and we used a Formlabs 3B+ printer. The bottom surface of the mold base was printed directly on the printer base without supports, to ensure a very flat surface. Both elastic and rigid resins were used, the former allowing an easier demolding but requires thicker features to ensure flatness. Specifically, we used Elastic 50A and High Temp resins from Formlabs.

After printing, molds were washed and cured according to manufacturer’s guidelines: bath in isopropanol under flow (in Form Wash, Formlabs) and 60 degrees celsius under UV light for 60 minutes in a curing station (Form Cure, Formlabs).

**Supp. Fig. 7.**
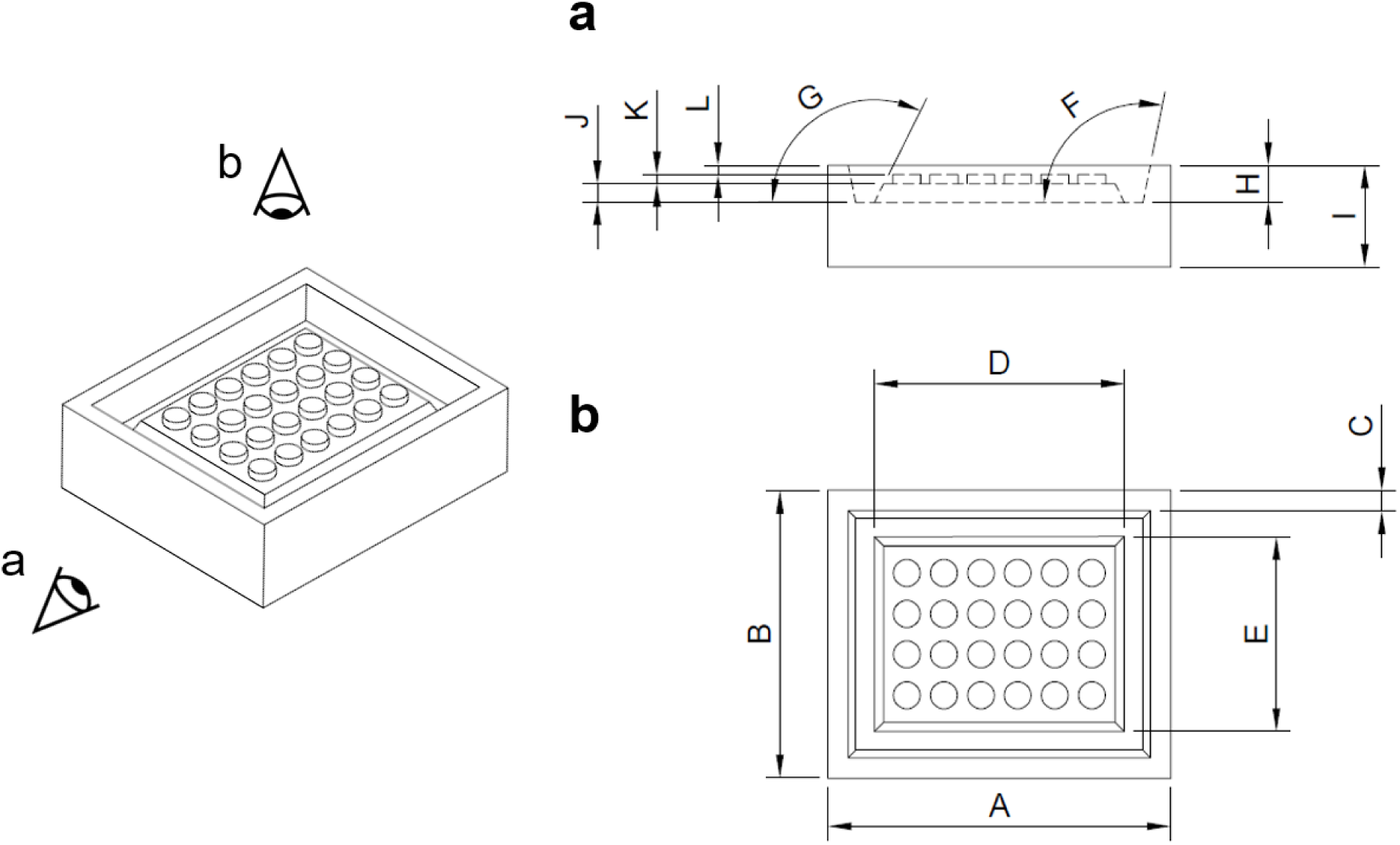
Technical drawing of representative mold (6×4F) with side and top views and critical dimensions.

## Sample Harvesting and Fixation

### Fixation & Harvest Method I: PFA based sample harvesting

#### Suitable for: (Soft) ECM-embedded culture models

1. Carefully aspirate the media from the organoid cultures.
2. Gently wash the organoid cultures with PBS and discard the PBS.
3. Add 4% PFA to the organoid domes and incubate at room temperature for 30 minutes. The hydrogel dome will dissociate in PFA.
4. Transfer the fixed organoids in PFA into a Falcon tube.
5. Gently pipette up and down along the side wall of the tube to dissolve any remaining hydrogel in the PFA.
6. Centrifuge the tube at 250xg for 5 minutes and carefully discard the supernatant containing PFA.
7. Wash the sample gently with PBS + 1% BSA and centrifuge the tube at 250xg for 5 minutes. Remove the supernatant.
8. Repeat the gentle wash with PBS + 1% BSA.
9. Remove the supernatant and gently resuspend the fixed organoid pellets in fresh PBS.
10. Store at 4°C until embedding.

### Fixation & Harvest Method II: Sample harvesting using Cell Recovery Solution

#### Suitable for: (Soft) ECM-embedded culture models

1. Carefully aspirate the media from the organoid cultures.
2. Carefully wash the sample with PBS without disturbing the dome.
3. Remove the wash buffer and add Cell Recovery Solution.
4. Place the plate in the fridge at 4°C for a minimum of 40 minutes.
5. Once the dome has softened or dissolved, resuspend the organoids in the solution by gentle pipetting. The dome will disintegrate, resulting in an organoid-suspension.
6. Use Advanced DMEM containing 1% BSA (to avoid organoid sticking to tip and tubes) to harvest organoids from the plate into an Eppendorf tube.
7. Centrifuge at 200xg for 5 minutes.
8. Carefully remove the supernatant, as the pellet is not very stable.
9. Wash the sample gently with cold PBS containing 1% BSA.
10. Centrifuge again and remove the supernatant.
11. Fix the organoids with 4% PFA for 30 minutes at room temperature.
12. After fixation, centrifuge the tube at 250xg for 5 minutes and carefully discard the supernatant containing PFA.
13. Wash the sample gently with PBS + 1% BSA and centrifuge the tube at 300xg for 5 minutes. Remove the supernatant.
14. Repeat the gentle wash with PBS + 1% BSA.
15. Remove the supernatant and gently resuspend the fixed organoid pellets in fresh PBS.
16. Store at 4°C until embedding.

### Fixation & Harvest Method III: Direct (in-situ) sample fixation

#### Suitable for: Single, mid-large sized models and tissue, stiff ECM-embedded models, transwell monolayer models

1. Carefully aspirate the media from the organoid cultures.
2. Carefully wash the sample once with PBS without disturbing the sample within the culture vessel.
3. Fix with 4% PFA overnight at 4°C or for 30-120 minutes at room temperature, based on the sample size within the culture vessel.
4. After the fixation, wash the sample three times with PBS. Discard the PBS wash.
5. Add fresh PBS to the sample.
6. Store at 4°C until embedding.

### Fixation & Harvest Method IV: Direct sample fixation of small organoids

#### Suitable for: Small suspension culture models

1. Transfer the organoids in media into an Eppendorf tube.
2. Centrifuge the tube at 300xg for 5 minutes and carefully aspirate the supernatant.
3. Gently wash the sample with PBS + 1% BSA and centrifuge the tube at 300xg for 5 minutes. Remove the supernatant.
4. Fix the sample with 4% PFA for 30 minutes at room temperature or overnight at 4°C.
5. Centrifuge the tube at 300xg for 5 minutes and carefully discard the supernatant containing PFA.
6. Wash the sample with PBS + 1% BSA and centrifuge the tube at 300xg for 5 minutes. Remove the supernatant.
7. Repeat the wash with PBS + 1% BSA.
8. Remove the supernatant and gently resuspend the fixed organoid pellets in fresh PBS.
9. Store at 4°C until embedding.

#### Preparation of histoarrays from histomolds

1. Select the appropriate histomold suited to the intrinsic and extrinsic factors described in Table 1 in conjunction with histomolds available (Suppl. Fig. 2).
2. Retrieve the Histogel™ from the fridge and thaw it in the microwave for approximately 10-20 seconds until it liquefies.

<u>Note</u>: Place the Histogel™ tube with a loosened cap inside a 50 mL Falcon tube with its lid slightly closed to prevent the Histogel™ from boiling over. Keep the Histogel™ at ∼ 64°C while working to maintain its liquid state.
3. Fill the histomold with Histogel™ by pipetting from the corners first, ensuring that you avoid creating bubbles and that the gel is evenly distributed on the surface. It is important to have the Histogel™ filled level on the surface, ensuring a flat surface once cast out of the mold.
4. Polymerize the Histogel™ in the mold for 20 minutes at 4°C in the fridge.
5. After polymerization, gently remove the Histogel™ from the mold and transfer it to a petri dish or Parafilm (non-sticky/hydrophobic surface) by using spatulas and squeezing it out. The resulting, cast-out Histogel™ is from here onward referred to as “histoarray” (see. Fig. 1a Schematic).

<u>Optional:</u> For added stability during sample transfer, place the trimmed histoarray into 24mm x 30mm disposable base mold.

### Optional: Labelling samples to ease identification of samples during sectioning

Identification of the depth positioning of fixed samples within an FFPE block during FFPE sectioning is critical for choosing the best possible FFPE slices for staining. For example, choosing sections too early or too far along during the sectioning process can lead to FFPE sections with suboptimal representation of samples or no biological material on the section. However, identification of appropriate cutting depths can be challenging for a variety of reasons: for explants, which vary slightly in size and buoyant properties (as discussed above), identification of fragments may be challenging due to poor colour contrast between wax and pale tissues, or due to fragment sizes much smaller than 1 mm^3^, as may be needed when testing numerous conditions from an initially scarce surgical resection or biopsy. This can be equally challenging when dealing with small spheroids or organoids liberated from ECM. These challenges are especially difficult for inexperienced users or users dealing with a novel type of sample in conjunction with histomolds. It is therefore advisable to improve the contrast between the CIVM being embedded and the surrounding material by staining fixed CIVMs with hematoxylin/eosin solution (Mayer’s hemalum solution), prior to embedding into histomolds, especially when beginning to use the histomolds. Towards this, fixed samples are ideally retained in well plates. A drop, equivalent to approx 5-10uL solution, is added to each well containing the sample in PBS. Samples are allowed to take up the dye at room temperature for 10-30 min, after which PBS + hematoxylin/eosin are removed from the wells using a small-bore tip (to ensure sample exclusion), washed once in PBS and then added to histomolds as described below. If samples are not in well plates, this process can be carried out on sheets of parafilm. Such pre-staining with hematoxylin/eosin greatly aids the identification of ideal sample planes within the FFPE block, as the colouring is retained all the way through FFPE sectioning. Importantly, this staining does not have repercussions on additional H&E staining, nor on (auto)fluorescence levels in mIF stainings.

### Sample transfer into histoarray and post-processing

The histomolds described within this manuscript were specifically designed for certain models, sizes and culture vessels in house. However, this is a non-exhaustive list and a constant, reiterative process of adjusting and optimizing the molds further is needed, as new models (might) demand slightly different sized wells or depths, for example. However, the same histomold often accommodates models of different organs of origins or culture methods and even sizes, depending on the user’s preference. The described embedding approaches can be adjusted to needs or preferences of the users and simply outline a variety of workflows we heuristically grouped into four main approaches covering most CIVMs.

In general, the samples can be handled using spatulas, tweezer, pipette tips, or a puncher depending on the model/application/matrix (see below). It is recommended to embed the samples in an asymmetric layout to easily identify samples post-processing and -sectioning, minimizing the chance of mislabeling (see Suppl. Fig. 6d). Furthermore, remove as much excess liquid carried over from the samples as possible before transferring into the histoarray to ensure homogenous dehydration results.

In cases where histomolds are to be filled completely, thereby yielding arrays with no evident orientation, it is further recommended to add an additional marker, in order to be able to unequivocally assign the correct orientation of FFPE sections: After filling and polymerizing the histomold completely, the top right corner of the histomold is cut away (see Suppl. Fig. 1A), the mold is placed within the histology cassette and an appropriately-sized piece of chicken breast tissue (previously fixed in 4% PFA and stored in PBS+sodium azide at 4 degrees) is added to the Histogel-free corner. The chicken breast is processed alongside the sample in all subsequent steps. Each FFPE section therefore contains a section of chicken breast, positioned in the top right corner of the CIVM histoarray. Chicken muscle is easily visible in H&E sections; in mIF stainings its autofluorescence can be utilized to orient the array to ease analysis. An alternative orientation control is a synthetic antigen gel made of bovine serum albumin (BSA) and containing pigment^89^. To prepare this type of orientation control, add 100 uL of tissue marking dye (TMD-GN, TBS) to 500 uL of 12.5% BSA in PBS (v/v) in a 1.5 mL Eppendorf tube, vortex the tube, heat at 85°C for 10 min on a Thermomixer (Eppendorf), remove the resulting solidified gel from the tube, and trim it into 2 mm x 2 mm x 5 mm pieces. These gel pieces can be placed directly into the notched, Histogel-free corner of the histoarray and processed alongside the sample, as described above, or the pieces can be wrapped in biopsy paper, processed separately in a tissue cassette, and incorporated at the time of paraffin embedding. These BSA gels do not come off the slide during antigen retrieval and are readily visualized on H&E sections, IHC, and IF.

### Transfer Method I: Single, mid-large sized models and tissues (see Fig. 2)

1. Remove the remaining liquid in the culture vessel and transfer the fixed sample onto parafilm using spatulas, tweezers or cut-off P1000 tips.
2. Aspirate any excess liquid carried over.
3. Place the samples into the histoarray using spatulas, tweezers or pieces of parafilm.
4. Repeat the previous step in case multiple samples are at hand.
5. Continue with step A below.

### Transfer Method II: Epithelial monolayer models on 24-well transwells (see Fig. 3)

1. Remove the fixed transwell samples from the plate.
2. Either use a scalpel or a biopsy puncher to remove the sample from the transwell insert, including the membrane.
3. Briefly place the transwell on parafilm (with the polymer facing downward) before transferring it into the histoarray.

<u>Note</u>: Do not push the transwell sample into the well, as this might lead to artifacts (e.g. rolling of the specimen at the bottom leading to undesired artifacts).
4. Repeat the previous step in case multiple samples are at hand.
5. Continue with step A below.

### Transfer Method II: Epithelial monolayer models on 96-well transwells (see Fig. 3)

1. Remove the fixed transwell samples from the plate.
2. Using forceps, grasp the edge of each transwell insert to peel off the sample with the transwell membrane.
3. Transfer the transwell directly into the histoarray.
4. Repeat the previous step for all samples.
5. Continue with step A below.

### Transfer Method III: Soft ECM-embedded culture models and small suspension culture models (see. Fig. 4)

1. Aspirate liquid in the Eppendorf tubes containing the fixed samples to ensure a dry organoid pellet (if necessary, centrifuge one more time).
2. Resuspend the samples with liquified Histogel™.

<u>Note</u>: The volume depends on the well size of the intended mold and amount of mold-wells you intend to fill with your samples. Ideally, the denser the better to enable a higher number of biological material per section.
3. Dispense organoid-Histogel™ suspension directly into the wells of the histoarray.
4. Repeat the previous step in case multiple samples are at hand.
5. Continue with step A below.

### Transfer Method IV: Stiff ECM-embedded (co-) culture models (see Fig. 5)

#### Stiff, collagen-based ECM preparation

To enable handling and embedding of samples without perturbing the spatial integrity, a stiff ECM is crucial. Recently, we’ve shown that a Rat Collagen I mixture with Matrigel (1:1 ratio, v/v) mixture is sufficient to retain the spatial integrity of the sample upon PFA fixation and transfer into the histoarray^83,84^. Besides the Rat Collagen I, we’ve also used Bovine Collagen (Telo-Col 6) with its commercially available neutralization solution that can be used to retain the integrity of the matrix (100% or mixed with Matrigel). Briefly, the stiff ECM was prepared following manufacturers ratios by resuspending 350 µL Telo-Col 6 with 40 µL of Neutralization Solution and mixed thoroughly (alternatively vortexed shortly). Next, the neutralized collagen was thoroughly mixed with 100 µL of Matrigel before centrifuging it for 30 sec at 20000 g at 4°C to avoid bubble formation.

Afterwards, samples can be resuspended with this hydrogel mixture and dispensed into a 24-well plate.

<u>Note</u>: The more hydrophobic the plate, the smoother the workflow described below (e.g. CytoOne CC7682-7524) as the dome will detach more easily. In addition, as different plates might lead to different sizes of domes (in respect to the total volume of the dome), the size of the dome (diameter) should be assessed once. The choice of puncher size and histomold-well diameter depends on this.

1. Remove any liquid from the wells containing the fixed samples.
2. Add 400 µL of Histogel™ to the sample-containing wells, submerging the sample fully.
3. Place the plate for a minimum of 25 min in the 4°C fridge to ensure full polymerization.
4. Afterwards, use a bent spatula and carefully scoop out the Histogel™-disk containing your sample and place it on parafilm.

i. Note: It can happen that the dome does not detach from the bottom of the plate with the Histogel™-disk. However, due to its rigid ECM, it can alternatively be directly removed from the plate using a spatula.
5. Use a puncher with the appropriate diameter for your dome size and histomold.
6. Punch the dome out of the Histogel™ card and discard the excess Histogel™.
7. Use two spatulas or parafilm pieces in order to transfer the dome into the histoarray.

<u>Note</u>: The diameter of the puncher and histomold-wells should be exactly the same as this facilitates perfect placement of the samples. For example, the cultures in Fig. 6 were cultured in 10 µL of matrix (see Fig. 6). The size of the dome was measured, the histomold wells specifically designed to accommodate the dome (max. 4.1 mm) and a 4 mm puncher selected to retrieve the samples and perfectly place them into the wells of the 5×4F flat bottom histomold (Suppl. Fig. 2).
8. Repeat steps 5-8 in case multiple samples are at hand.
9. Continue with step A below.

### From here, applicable to all transfer-workflows described above

A. Allow the samples to dry in the histoarray for 2 minutes.
B. <u>Critical</u>: Before completely filling with histoarray, "glue" the samples onto the histoarray by slowly dispensing a few microliters of Histogel™ around each sample using a P10 pipette. Wait 5 minutes to allow for polymerization of the Histogel™.

<u>Note</u>: Filling Histogel™ directly on top may cause the samples to lift from the histoarray wells (buoyancy), especially in the case of small/light samples. Carefully pre-polymerizing the samples within the wells before filling the rest of the histoarray prevents this.
<u>Note:</u> Not necessary for ‘Soft-embedded culture models’, which are dispensed in Histogel™ directly into the histoarray (see above).
C. Slowly fill up the rest of the mold with Histogel™, keeping it level.

a. <u>Note</u>: for 96-well transwell samples, add just enough Histogel to stabilize the samples and prevent them from folding over. Adding a large volume of Histogel to fully encapsulate the samples will increase the thickness of the histoarray and could increase the risk of distortion artifacts during processing.
D. Place the Histogel™-covered histoarrays in the 4°C fridge for 10-15 minutes to polymerize the Histogel™.
E. Store any remaining Histogel™ at 4°C.
F. Once the Histogel™ is polymerized, trim the excess corners of the samples embedded in Histogel™ if needed, without disturbing the lanes containing samples, to fit them in the cassette.
G. Place the sample-embedded histoarray inside a histo-cassette without flipping the histoarray.

<u>Critical:</u> As the true flat surface is the bottom of the cast histoarray and therefore enables sectioning through all samples at once, the histoarray containing the samples should not be flipped around during the FFPE-embedding process.
Note: It can help to cut a corner of the histoarray in order to identify the orientation later more easily.
H. Place the cassette in 10% formalin for 4 hours at RT and proceed with sample dehydration and paraffin embedding afterwards.

<u>Note:</u> Store thick Transwell histoarrays for 24 hours in 10% formalin at RT followed by 24 hours in 70% ethanol to help reduce processing artifacts.

### Sample Dehydration, Paraffin Embedding and Sectioning

The sample processing is performed in a fully automated tissue processor HistoCore PEARL (Leica).

1. Place the basket with cassettes in the chamber, select the appropriate program, and start the machine.
2. Different types of sample blocks require different processing lengths depending on their size or thickness. Refer to the table below for examples. We recommend longer dehydration times for thicker blocks.
3. The processing retort is initially filled with reagent from the first reaction bottle.
4. After the set time, the reagent is returned to its respective bottle, and the processing retort is filled with reagent from the next bottle. This process continues sequentially.
5. Thirteen reagents are exchanged in one cycle within the processing retort.
6. After the processing cycle is complete, remove the cassette basket and proceed with embedding.

**Table 3:**
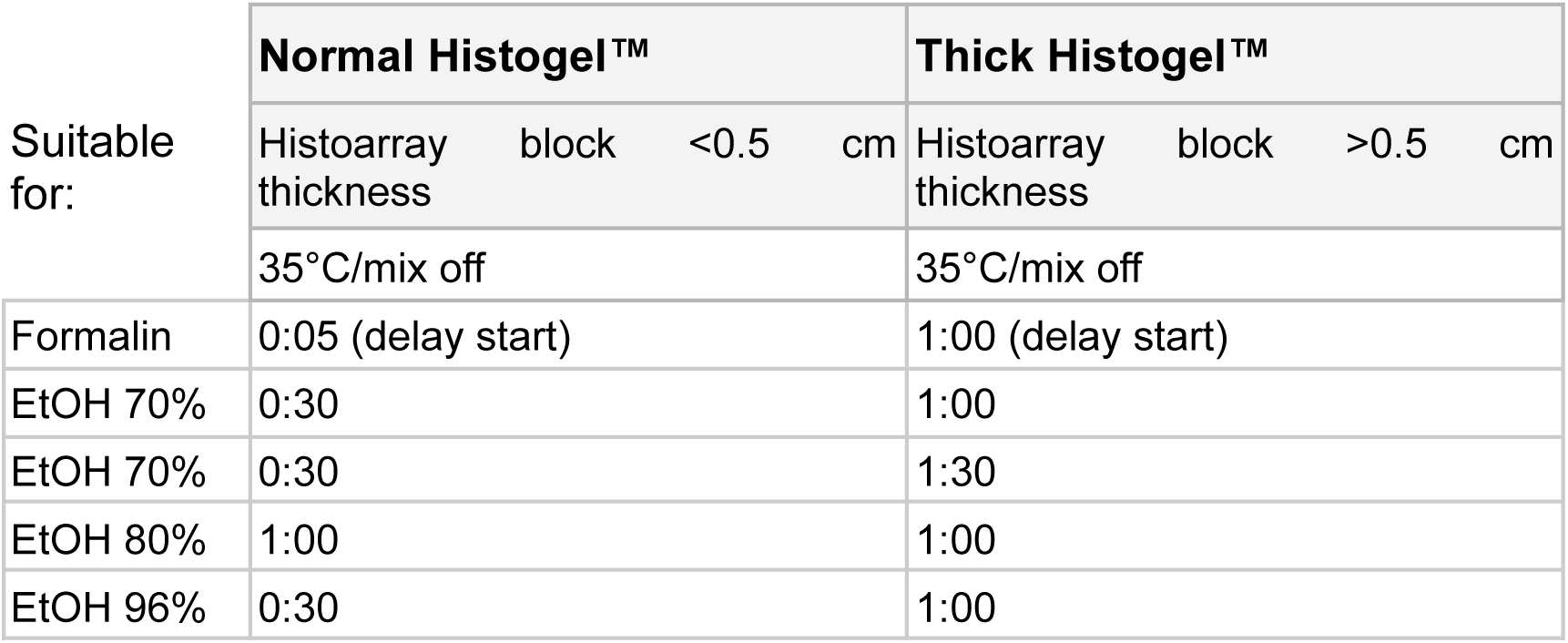
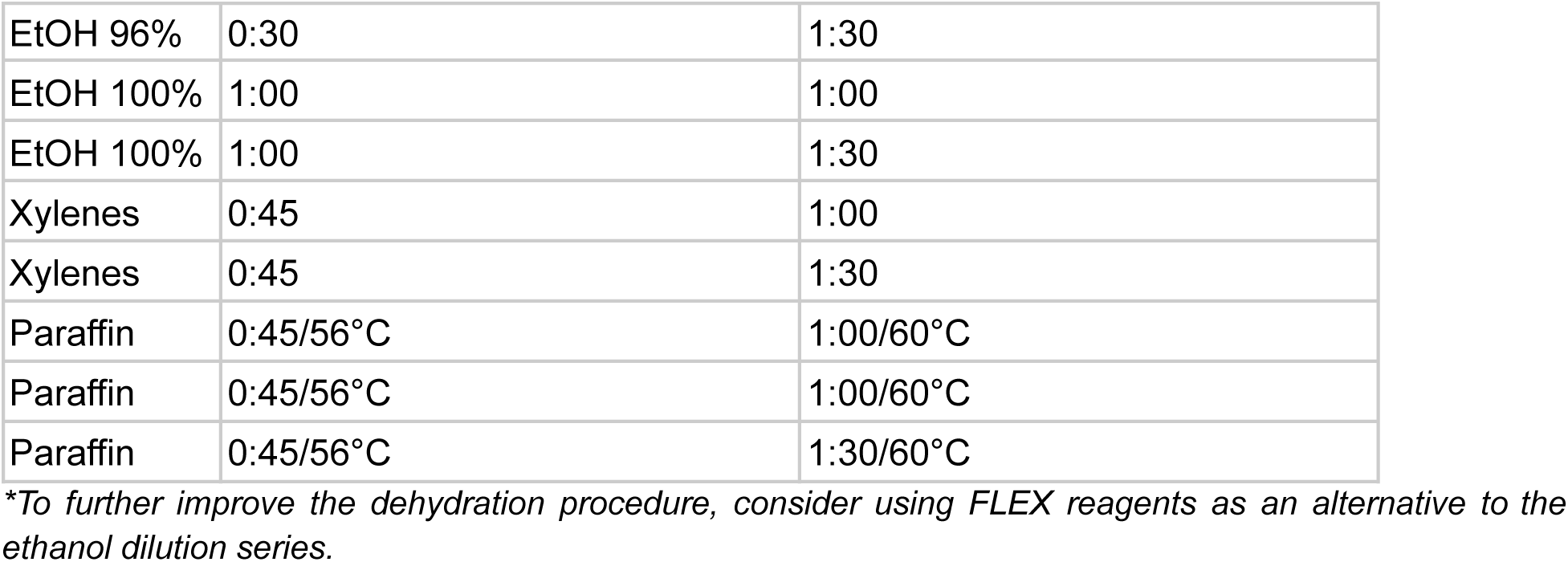
Table of Specified Programs for Processing HistoGel Blocks in the HistoCore PEARL Tissue Processor.

### Paraffin embedding

Embedding was performed on a Medite Embedding station TES99.

1. Take an appropriate metallic-mold and fill half of it.
2. Open the cassette containing the processed histoarray. Carefully transfer the histoarray from the cassette to the metallic mold filled with paraffin.

<u>Critical:</u> As described above, do not flip the histoarray as the bottom side is intended to be on the bottom of the metallic mold to ensure correct orientation for subsequent sectioning.
3. Transfer the metallic mold containing the processed histoarray to the cold section of the plate. Gently press the histoarray to the bottom of the mold with tweezers.

<u>Critical:</u> Make sure to equally apply pressure, ensuring the bottom of the histoarray is in one plane.
4. Cover the mold with the corresponding cassette and fill it with liquid paraffin up to the edge of the cassette. Place it on the cold plate until properly solidified.
5. After cooling, separate the paraffin block from the mold, remove any excess paraffin, and proceed with sectioning.

### Microtome sectioning

Sectioning was performed on a Thermo Microm HM355S.

1. Fill the water bath with distilled water and adjust the temperature to 42°C.
2. Trim FFPE-blocks at a thickness of 10 to 20 µm at room temperature until the first samples appear, then place the FFPE-blocks on a cold plate or an ice bath for 10-15 min, which hydrates and cools the blocks, allowing for better ribbon formation.

<u>Note:</u> The Histogel™ has a distinct appearance in contrast to the paraffin, as do the CIVMs in the sections. A trained person will detect even small organoids by eye with ease, whereas non-trained personnel could use the optional labelling step to identify the models during sectioning (described above).
3. Once the FFPE-blocks are sufficiently cold, section the FFPE-blocks at a thickness of 3-4 µm or thicker sections, depending on the method or requirement. The sections should meet the following criteria:

i. All embedded samples must be captured in one plane.

<u>Note:</u> If all tips above have been followed, this should be the case.
ii. The section must cover the largest surface area of the embedded tissue.

<u>Note</u>: Once all samples are captured in one section, perform several consecutive sections at different depths in order to evaluate the section-depth (by H&E, see below).
iii. The section must not be wrinkled or folded, this might lead to unwanted artifacts during imaging.
4. Cut serial sections of 6 to 7 sections per piece and place them on the surface of the water bath.

<u>Critical</u>: For transwell histoarrays, we recommend setting the water bath temperature to 36°C and collecting only 1-2 sections at a time, optimally within 5-10 seconds for best results. Flotation times of up to 60 seconds can be used with relatively minor drop off in section quality. However, if using a standard water bath temperature (41-42°C), flotation times longer than 5-10 seconds result in significant sample waviness and increased sample loss. If a ribbon of serial sections is needed, collect the appropriate number of sections onto the microtome blade holder and transfer to a water bath in smaller ribbons of 2-3 sections.

<u>Note:</u> Separate individual sections using forceps cooled on an ice bath to prevent the transwell samples within the paraffin section from becoming stretched.
5. Transfer the sections from the water bath onto Superfrost or Gold Superfrost slides.

<u>Note</u>: Use Gold Superfrost slides for thin, long samples (e.g. transwells) to avoid lifting during processing.
6. After sectioning, place the slides in an incubator at 37°C overnight.
7. The slides can be stored at room temperature for up to one year.

Depending on the CIVM, we have a two-way sectioning approach.

a) In case of large samples (e.g. explants or assembloids), we recommend obtaining sections at multiple depths to cover the heterogeneity of the sample.
b) In case of small samples (e.g. small organoid domes, small kidney organoids), we recommend obtaining as many consecutive sections as possible without trimming at different depths or re-trimming when intending to section the FFPE-block another day to avoid sample loss.

## Histology techniques

### Hematoxylin and Eosin (H&E) Staining

H&E staining was executed in a fully automated manner following the standard protocol on the Ventana HE600 stainer (Roche Tissue Diagnostics).

### Alcian Blue Staining

AlcianBlue (5279194001, Ventana) -PAS staining (5279291001, Ventana) was performed fully automated on the Ventana Benchmark (Roche Tissue Diagnostics). Briefly, after three cycles of deparaffinization, slides were washed and 200 µL of PAS Alcian blue incubated for 16 min at 37°C. Wash was repeated, and 200 µL of Pas periodic acid was applied for 4 min, before another wash and 12 min of PAS Schiffs staining. Next, slides were washed again and 200 µL PAS neutralizer was added and incubated for 4 min. Lastly, slides were washed with 100 µL before counterstaining the slide with 200 µL of PAS hematoxylin for 8 min. Afterwards, slides were coverslipped as indicated in the manufacturer’s instructions.

### Immunohistochemistry (IHC)

Stainings were performed using Ventana Discovery Ultra automated tissue stainer (Roche Tissue Diagnostics, Tucson AZ USA). Primary antibodies and concentrations were validated in establishment IHC runs (except for pre-diluted Ventana primary antibodies), prepared in Discovery Ab diluent and then subjected to a dispenser for automated application. Specific incubation times might change between antibodies.

1. Bake the slides first at 60°C for 8 min and subsequently further heat up to 69°C for 8 min for subsequent deparaffinization. Repeat this cycle three times.
2. Perform heat-induced antigen retrieval with Tris-EDTA buffer pH 7.8 at 95°C for a total of 40 min.
3. Apply Discovery Goat Ig Block and incubate for 32 min.
4. Apply Discovery Inhibitor for 16 min.
5. Apply primary antibodies for 40 min.
6. Detect primary antibodies using anti-species secondary antibodies conjugated to horseradish peroxidase (HRP) for 16 min and subsequently visualize by conversion of 3,3’-Diaminobenzidine (DAB).
7. Counterstain specimens with Hematoxylin and Bluing Reagent.
8. Dehydrate with a standard series of alcohol (70% v/v, 96% v/v, 100% v/v, 100% v/v) and Xylol baths (100% v/v), then mount slides fully automated using the RCM7000 coverslipper and a standard histoglue.
9. Dry the slides for at least 2 h prior to imaging.

### Multiplexed Immunofluorescence (mIF; opal dyes)

Prepare Opal dyes as described by the manufacturer instructions. Stainings were performed using Ventana Discovery ultra automated tissue stainer (Roche Tissue Diagnostics, Tucson AZ USA). Primary antibodies and concentrations were validated in establishment IHC runs (except for pre-diluted Ventana primary antibodies), prepared in Discovery Ab diluent and then subjected to a dispenser for automated application. The order of the primary antibodies and corresponding dyes was further determined during establishment runs. Specific incubation times might change between antibodies.

1. Bake the slides first at 60°C for 8 min and subsequently further heat up to 69°C for 8 min for subsequent deparaffinization. Repeat this cycle three times.
2. Perform heat-induced antigen retrieval with Tris-EDTA buffer pH 7.8 at 95°C for a total of 40 min.
3. Apply Discovery Goat Ig Block and incubate for 32 min.
4. Apply Discovery Inhibitor for 16 min.
5. Apply primary antibodies for 40 min.
6. Apply the respective anti-species secondary antibodies conjugated to horseradish peroxidase (HRP) for 16 min.
7. Subsequently, apply respective Opal dye (starting with 480 to 780) and incubate for 12 min at 37°C.
8. After every application of a primary antibody, respective secondary antibody and Opal dye, an antibody neutralization and denaturation step (4 min at 92°C) is applied to remove residual antibodies and HRP, before starting the staining cycle again with the Discovery Inhibitor blocking step (Step 4).
9. In the seven sequence, after the primary an additional Goat IgG blocking step is performed for 16 min, followed by the secondary antibody-HRP. Next, Opal TSA reagent is applied, followed by the Opal dye 780.
10. Slides are manually mounted using ProLong™ Gold Antifade Mountant. Slides are dried for at least 2 hours prior to imaging.

### Sample preparation for MERFISH spatial transcriptomics

#### Histomold embedding

Human intestinal organoids growing in domes were washed with PBS and directly fixed with 4% PFA in nuclease-free PBS for 30 min at 4°C. After fixation, the organoids were carefully resuspended by pipetting, transferred to a 50 ml Falcon tube and spun at 300g, 4°C for 4 min. The pellet was washed once with cold nuclease-free PBS with 1% BSA, resuspended in cold nuclease-free PBS with 1% BSA, transferred in a 2 ml tube and spun at 300g, 4°C for 4 min. For embedding, the supernatant was removed and the organoids were transferred into 8 wells of a flat-bottom (5×4F) histomold. After drying for ∼2-3 min, a small drop of histogel was added on top of each well. After ∼1 min, the gel was filled up and allowed to solidify at 4°C for a few minutes. The histogel was placed in a histology cassette and fixed with 10% formalin (HT501320, Sigma) for 1h at 4°C. The cassette was transferred into cold nuclease-free PBS until placed in the Leica for dehydration and paraffin embedding.

#### RNA isolation and DV_200_

The FFPE block was sectioned with a Microtome under RNAse-free conditions. Two sections of 10 μm were deparaffinized with the Deparaffinization solution (19093, Qiagen) and used for RNA isolation with the Qiagen RNeasy FFPE kit (73504, Qiagen). DV_200_ was obtained by running the isolated RNA on an Agilent RNA 6000 Pico Kit (5067-1513) in the Agilent 2100 Bioanalyzer.

#### MERFISH spatial transcriptomics

The FFPE block was sectioned with a Microtome in RNAse-free conditions. A 4 μm section was placed on a Merscope FFPE Slide (10500102, Vizgen) and processed for MERFISH with Merscope FFPE Sample Preparation reagents (10400114, Vizgen) and user guide Formalin-Fixed Paraffin-Embedded Tissue Sample Preparation 91600112 Rev C (Vizgen). After anchoring pretreatment, the sample was stained with Cell Boundary Staining (10400118, Vizgen) followed by RNA anchoring and gel embedding. The sample was treated with Digestion Mix for 2h at 37°C before undergoing clearing for 24h at 47°C, autofluorescence quenching and further clearing for 24h at 37°C. A custom panel with 409 probes against duodenal cell marker genes was hybridized for ∼46h at 37°C. The slide was washed and imaged on the Merscope instrument (10000001, Vizgen) with a gene imaging kit (10400006, Vizgen) according to Vizgen user guide 91600001. Images were taken with Merscope Vizualizer software.

## Imaging

### IHC, H&E and Special Stains

HE, IHC and special stained slides were imaged with a brightfield whole-slide scanner at 40X (Hamamatsu, NanoZoomer S360).

### mIF with Opal Kits

#### Imaging with Vectra Polaris

1. Digitize mIF stainings using the Opal dyes from Akoya with multispectral imaging by the Vectra® Polaris™ (PerkinElmer) using the MOTiF™ technology at 20x magnification for all 7 colors (Opal 480, Opal 520, Opal 570, Opal 620, Opal 690 and Opal 780).
2. Adjust the laser exposure and intensity settings on multiple slides per staining panel. Scan the slides in a batch manner to ensure same imaging settings and cross-comparability for later image analysis with the image analysis software of choice.
3. Next, perform unmixing of the channels and tiling of the images with PhenoChart (v1.0.12) and inForm (v2.4).
4. The raw images were saved as .qptiff and later fused in HALO (Indica labs, v3.6.4134.396).

### Image Analysis

Image analysis of IHC and mIF images was performed with HALO (Indica Labs, v3.6.4134.396) and HALO AI (Indica Labs, v3.6.4134). Sample annotation was swiftly performed using the TMA module in HALO depending on the histomold utilized.

For single organoid detection and annotation (for both IHC and mIF images), a RandomForest (v2) classifier was used to distinguish organoids from background (e.g. matrix; minimum object size > 1000 µm). Following quick manual validation, organoids were handled as individual regions of interest (ROI) and cell quantification executed per ROI. The RandomForest (v2) classifier was integrated in the Area Quantification (v2.4.3) and Area Quantification FL (v.2.3.4) modules and used to quantify positive marker staining against either overall size of each individual object or normalized to the DAPI^+^ area of the object (as indicated in legend texts). The HighPlex FL (v4.2.14) module was used to perform nuclear segmentation based on DAPI^+^ cells, assisted by HALO AI’s integrated ‘AI default nuclear segmentation type’). Specific cell phenotypes were determined by co-localization of markers of interest with the DAPI^+^ nuclei (taking the cytoplasm radius (1.25 µm) and nuclear signals into account). By integrating the RandomForest classifier in the HighPlex module, e.g. localization of the T cells in the matrix or epithelium were determined and normalized to the tumoroid area. Secondary-only staining controls on a consecutive section or primary tissue served as a negative signal threshold to prevent biased adjustments. For some samples, negative controls were co-embedded within the same histoarray to ease unbiased threshold adjustment. For the analysis of antigen-expressing areas in explants, Area Quantification FL (v.2.3.4) was utilized to set the threshold for signal above background for EpCam and Ceacam5 stainings, in order to quantify the area of positivity relative to the area of cohesive explant tissue. Note that for explants generating substantial amounts of debris and loss of tissue cohesion during culture (such as explants generated from normal colon), DAPI signal was used to identify areas of cellular cohesive explant tissue, which were defined with manual ROIs. This effectively removed areas of debris and allowed for more accurate antigen quantification.

## Author contributions

M.F.H. conceived the study. M.F.H. wrote the manuscript with support from E.D.A., E.S., I.P. and N.G.. M.F.H., B.L., A.M.F., C.H. and E.S. optimized the histo-workflow. C.H. and E.S. co-developed and optimized the 96-well histomolds and protocol together with R.H. and L.N with support from I.P.. E.K. co-developed the 24-well histomold with I.P., J.A. and M.F.H.. B.L. and B.S. accumulated protocols with support from M.F.H.. R.O., M.F.H., J.A., I.P. and J.L.G.C. conceptualized, designed and optimized the histomolds with support of colleagues. E.D.A., J.A., E.K., J.S.D., R.L.S., C.H., A.M.F., G.B., E.S. contributed models and respective data, with support from G.C..

## Acknowledgements

We would like to express our gratitude to the entire Pathology and Applied Safety Sciences team at Roche pRED for providing access to their lab and instruments. Additionally, we wish to thank the following individuals and core labs at Genentech: Linda Rangell for project support and guidance; Emily Mu for timely generation of transwell samples to enable optimization of the transwell histomold design; and the Digital Pathology Image Analysis group in Research Pathology for whole slide scanning support. Furthermore, we would like to thank Davit Suter for sharing the murine ESC line SBR.

## Bibliography

1. Tang, X.-Y. et al. Human organoids in basic research and clinical applications. Signal Transduct. Target. Ther. 7, 168 (2022).

2. Clevers, H. Modeling Development and Disease with Organoids. Cell 165, 1586–1597 (2016).

3. Zhao, Z., et al. Organoids. Nat. Rev. Methods Prim. 2, 94 (2022).

4. Corrò, C., Novellasdemunt, L. & Li, V. S. W. A brief history of organoids. Am. J. Physiol.-Cell Physiol. 319, C151–C165 (2020).

5. Simian, M. & Bissell, M. J. Organoids: A historical perspective of thinking in three dimensions. J. Cell Biol. 216, 31–40 (2017).

6. Drost, J. & Clevers, H. Organoids in cancer research. Nat. Rev. Cancer 18, 407–418 (2018).

7. Fatehullah, A., Tan, S. H. & Barker, N. Organoids as an in vitro model of human development and disease. Nat. Cell Biol. 18, 246–254 (2016).

8. LeSavage, B. L., Suhar, R. A., Broguiere, N., Lutolf, M. P. & Heilshorn, S. C. Next-generation cancer organoids. Nat. Mater. 21, 143–159 (2022).

9. Hofer, M. & Lutolf, M. P. Engineering organoids. Nat. Rev. Mater. 6, 402–420 (2021).

10. Kim, J., Koo, B.-K. & Knoblich, J. A. Human organoids: model systems for human biology and medicine. Nat. Rev. Mol. Cell Biol. 21, 571–584 (2020).

11. Hughes, D. L., Hughes, A., Soonawalla, Z., Mukherjee, S. & O’Neill, E. Dynamic Physiological Culture of Ex Vivo Human Tissue: A Systematic Review. Cancers 13, 2870 (2021).

12. Dorrigiv, D., et al. Microdissected Tissue vs. Tissue Slices—A Comparative Study of Tumor Explant Models Cultured On-Chip and Off-Chip. Cancers 13, 4208 (2021).

13. Voabil, P. et al. An ex vivo tumor fragment platform to dissect response to PD-1 blockade in cancer. Nat. Med. 27, 1250–1261 (2021).

14. McAleer, C. W. et al. Multi-organ system for the evaluation of efficacy and off-target toxicity of anticancer therapeutics. Sci. Transl. Med. 11, (2019).

15. Vernetti, L. et al. Functional Coupling of Human Microphysiology Systems: Intestine, Liver, Kidney Proximal Tubule, Blood-Brain Barrier and Skeletal Muscle. Sci. Rep. 7, 42296 (2017).

16. Kim, H. J., Huh, D., Hamilton, G. & Ingber, D. E. Human gut-on-a-chip inhabited by microbial flora that experiences intestinal peristalsis-like motions and flow. Lab a Chip 12, 2165–2174 (2012).

17. Huh, D. et al. Reconstituting Organ-Level Lung Functions on a Chip. Science 328, 1662–1668 (2010).

18. Cingolani, E. et al. In vitro investigation on the impact of airway mucus on drug dissolution and absorption at the air-epithelium interface in the lungs. Eur. J. Pharm. Biopharm. 141, 210–220 (2019).

19. Stone, N. L., England, T. J. & O’Sullivan, S. E. A Novel Transwell Blood Brain Barrier Model Using Primary Human Cells. Front. Cell. Neurosci. 13, 230 (2019).

20. Wright, C. W. et al. Establishment of a 96-well transwell system using primary human gut organoids to capture multiple quantitative pathway readouts. Sci. Rep. 13, 16357 (2023).

21. Bluhmki, T. et al. Development of a miniaturized 96-Transwell air–liquid interface human small airway epithelial model. Sci. Rep. 10, 13022 (2020).

22. Dye, B. R. et al. In vitro generation of human pluripotent stem cell derived lung organoids. eLife 4, e05098 (2015).

23. Mun, S. J. et al. Generation of expandable human pluripotent stem cell-derived hepatocyte-like liver organoids. J. Hepatol. 71, 970–985 (2019).

24. Takasato, M. et al. Kidney organoids from human iPS cells contain multiple lineages and model human nephrogenesis. Nature 526, 564–568 (2015).

25. Sato, T. et al. Long-term Expansion of Epithelial Organoids From Human Colon, Adenoma, Adenocarcinoma, and Barrett’s Epithelium. Gastroenterology 141, 1762–1772 (2011).

26. Lancaster, M. A. et al. Cerebral organoids model human brain development and microcephaly. Nature 501, 373–379 (2013).

27. McCracken, K. W. et al. Modelling human development and disease in pluripotent stem-cell-derived gastric organoids. Nature 516, 400–404 (2014).

28. Lindemans, C. A. et al. Interleukin-22 promotes intestinal-stem-cell-mediated epithelial regeneration. Nature 528, 560–564 (2015).

29. Schreurs, R. R. C. E. et al. Human Fetal TNF-α-Cytokine-Producing CD4+ Effector Memory T Cells Promote Intestinal Development and Mediate Inflammation Early in Life. Immunity 50, 462–476.e8 (2019).

30. Holokai, L. et al. Increased Programmed Death-Ligand 1 is an Early Epithelial Cell Response to Helicobacter pylori Infection. PLoS Pathog. 15, e1007468 (2019).

31. Rogoz, A., Reis, B. S., Karssemeijer, R. A. & Mucida, D. A 3-D enteroid-based model to study T-cell and epithelial cell interaction. J. Immunol. Methods 421, 89–95 (2015).

32. Tsuruta, S. et al. Development of Human Gut Organoids With Resident Tissue Macrophages as a Model of Intestinal Immune Responses. Cell. Mol. Gastroenterol. Hepatol. 14, 726–729.e5 (2022).

33. Kang, S. et al. Complex in vitro models positioned for impact to drug testing in pharma: a review. Biofabrication 16, 042006 (2024).

34. Stokar-Regenscheit, N. et al. Complex In Vitro Model Characterization for Context of Use in Toxicologic Pathology: Use Cases by Collaborative Teams of Biologists, Bioengineers, and Pathologists. Toxicol. Pathol. 52, 123–137 (2024).

35. Stresser, D. M. et al. Towards in vitro models for reducing or replacing the use of animals in drug testing. Nat. Biomed. Eng. 8, 930–935 (2024).

36. Olsen, Timothy. R., et al. Processing cellular spheroids for histological examination. J. Histotechnol. 37, 138–142 (2014).

37. Guyon, J. & Daubon, T. Histological analysis of invasive glioblastoma organoids embedded in a 3D collagen matrix. STAR Protoc. 4, 102521 (2023).

38. Zhang, S. et al. An efficient and user-friendly method for cytohistological analysis of organoids. J. Tissue Eng. Regen. Med. 15, 1012–1022 (2021).

39. Havnar, C. et al. Histogel-based techniques for embedding organoids in paraffin blocks enable high throughput downstream histopathological analyses. J. Histotechnol. ahead-of-print, 1–12 (2024).

40. Beachley, V. Z. et al. Tissue matrix arrays for high-throughput screening and systems analysis of cell function. Nat. Methods 12, 1197–1204 (2015).

41. Kononen, J. et al. Tissue microarrays for high-throughput molecular profiling of tumor specimens. Nat. Med. 4, 844–847 (1998).

42. Yan, P., Seelentag, W., Bachmann, A. & Bosman, F. T. An Agarose Matrix Facilitates Sectioning of Tissue Microarray Blocks. J. Histochem. Cytochem. 55, 21–24 (2006).

43. Thomsen, A. R. et al. A deep conical agarose microwell array for adhesion independent three-dimensional cell culture and dynamic volume measurement. Lab a Chip 18, 179–189 (2017).

44. Simeone, K. et al. Paraffin-embedding lithography and micro-dissected tissue micro-arrays: tools for biological and pharmacological analysis of ex vivo solid tumors. Lab a Chip 19, 693–705 (2019).

45. Kabadi, P. K. et al. Into the depths: Techniques for in vitro three-dimensional microtissue visualization. BioTechniques 59, 279–286 (2015).

46. Gabriel, J., Brennan, D., Elisseeff, J. H. & Beachley, V. Microarray Embedding/Sectioning for Parallel Analysis of 3D Cell Spheroids. Sci. Rep. 9, 16287 (2019).

47. Chen, J. et al. Cerebral Organoid Arrays for Batch Phenotypic Analysis in Sections and Three Dimensions. Int. J. Mol. Sci. 24, 13903 (2023).

48. Heub, S. et al. Coplanar embedding of multiple 3D cell models in hydrogel towards high-throughput micro-histology. Sci. Rep. 12, 9991 (2022).

49. Powley, I. R. et al. Patient-derived explants (PDEs) as a powerful preclinical platform for anti-cancer drug and biomarker discovery. Br. J. Cancer 122, 735–744 (2020).

50. Karekla, E. et al. Ex Vivo Explant Cultures of Non–Small Cell Lung Carcinoma Enable Evaluation of Primary Tumor Responses to Anticancer Therapy. Cancer Res. 77, 2029–2039 (2017).

51. Gerlach, M. M. et al. Slice cultures from head and neck squamous cell carcinoma: a novel test system for drug susceptibility and mechanisms of resistance. Br. J. Cancer 110, 479–488 (2014).

52. Naipal, K. A. T. et al. Tumor slice culture system to assess drug response of primary breast cancer. BMC Cancer 16, 78 (2016).

53. Demetriou, C. et al. An optimised patient-derived explant platform for breast cancer reflects clinical responses to chemotherapy and antibody-directed therapy. Sci. Rep. 14, 12833 (2024).

54. Gavert, N. et al. Ex vivo organotypic cultures for synergistic therapy prioritization identify patient-specific responses to combined MEK and Src inhibition in colorectal cancer. Nat. Cancer 3, 219–231 (2022).

55. Runge, A. et al. Patient-derived head and neck tumor slice cultures: a versatile tool to study oncolytic virus action. Sci. Rep. 12, 15334 (2022).

56. Lang, N. J. et al. Ex vivo tissue perturbations coupled to single-cell RNA-seq reveal multilineage cell circuit dynamics in human lung fibrogenesis. Sci. Transl. Med. 15, eadh0908 (2023).

57. Ansari, M. Y. et al. Mitochondrial dysfunction triggers a catabolic response in chondrocytes via ROS-mediated activation of the JNK/AP1 pathway. J. Cell Sci. 133, jcs247353 (2020).

58. Martínez-Sabadell, A. et al. The target antigen determines the mechanism of acquired resistance to T cell-based therapies. Cell Rep. 41, 111430 (2022).

59. Baldassi, D., Gabold, B. & Merkel, O. M. Air−Liquid Interface Cultures of the Healthy and Diseased Human Respiratory Tract: Promises, Challenges, and Future Directions. Adv. NanoBiomed Res. 1, (2021).

60. Swart, A. L. et al. Pseudomonas aeruginosa breaches respiratory epithelia through goblet cell invasion in a microtissue model. Nat. Microbiol. 9, 1725–1737 (2024).

61. Lee, R. E. et al. Air-Liquid interface cultures to model drug delivery through the mucociliary epithelial barrier. Adv. Drug Deliv. Rev. 198, 114866 (2023).

62. Leach, T. et al. Development of a novel air–liquid interface airway tissue equivalent model for in vitro respiratory modeling studies. Sci. Rep. 13, 10137 (2023).

63. Gras, D. et al. Epithelial ciliated beating cells essential for ex vivo ALI culture growth. BMC Pulm. Med. 17, 80 (2017).

64. Hiemstra, P. S., Tetley, T. D. & Janes, S. M. Airway and alveolar epithelial cells in culture. Eur. Respir. J. 54, 1900742 (2019).

65. Sato, T. et al. Single Lgr5 stem cells build crypt-villus structures in vitro without a mesenchymal niche. Nature 459, 262–265 (2009).

66. Co, J. Y., Klein, J. A., Kang, S. & Homan, K. A. Suspended hydrogel culture as a method to scale up intestinal organoids. Sci. Rep. 13, 10412 (2023).

67. Bacac, M. et al. A Novel Carcinoembryonic Antigen T-Cell Bispecific Antibody (CEA TCB) for the Treatment of Solid Tumors. Clin. Cancer Res. 22, 3286–3297 (2016).

68. Tiernan, J. P. et al. Carcinoembryonic antigen is the preferred biomarker for in vivo colorectal cancer targeting. Br. J. Cancer 108, 662–667 (2013).

69. Campos-da-Paz, M., Dórea, J. G., Galdino, A. S., Lacava, Z. G. M. & Santos, M. de F. M. A. Carcinoembryonic Antigen (CEA) and Hepatic Metastasis in Colorectal Cancer: Update on Biomarker for Clinical and Biotechnological Approaches. Recent Pat. Biotechnol. 12, 269–279 (2018).

70. Fujii, M. et al. Human Intestinal Organoids Maintain Self-Renewal Capacity and Cellular Diversity in Niche-Inspired Culture Condition. Cell Stem Cell 23, 787–793.e6 (2018).

71. Zhang, Y. & Zhang, Z. The history and advances in cancer immunotherapy: understanding the characteristics of tumor-infiltrating immune cells and their therapeutic implications. Cell. Mol. Immunol. 17, 807–821 (2020).

72. Mellman, I., Coukos, G. & Dranoff, G. Cancer immunotherapy comes of age. Nature 480, 480–489 (2011).

73. Wang, Y., Wang, M., Wu, H. & Xu, R. Advancing to the era of cancer immunotherapy. Cancer Commun. 41, 803–829 (2021).

74. Wang, Q. et al. Role of tumor microenvironment in cancer progression and therapeutic strategy. Cancer Med. 12, 11149–11165 (2023).

75. Arner, E. N. & Rathmell, J. C. Metabolic programming and immune suppression in the tumor microenvironment. Cancer Cell 41, 421–433 (2023).

76. Visser, K. E. de & Joyce, J. A. The evolving tumor microenvironment: From cancer initiation to metastatic outgrowth. Cancer Cell 41, 374–403 (2023).

77. Dekkers, J. F. et al. Uncovering the mode of action of engineered T cells in patient cancer organoids. Nat. Biotechnol. 41, 60–69 (2023).

78. Zhong, Z. et al. Human immune organoids to decode B cell response in healthy donors and patients with lymphoma. Nat. Mater. 1–15 (2024) doi:10.1038/s41563-024-02037-1.

79. Esposito, A. et al. Colorectal cancer patients-derived immunity-organoid platform unveils cancer-specific tissue markers associated with immunotherapy resistance. Cell Death Dis. 15, 878 (2024).

80. Zhou, G. et al. Modelling immune cytotoxicity for cholangiocarcinoma with tumour-derived organoids and effector T cells. Br. J. Cancer 127, 649–660 (2022).

81. Kroll, K. T. et al. Immune-infiltrated kidney organoid-on-chip model for assessing T cell bispecific antibodies. Proc. Natl. Acad. Sci. 120, e2305322120 (2023).

82. Walsh, L. A. & Quail, D. F. Decoding the tumor microenvironment with spatial technologies. Nat. Immunol. 24, 1982–1993 (2023).

83. Harter, M. F. et al. Analysis of off-tumour toxicities of T-cell-engaging bispecific antibodies via donor-matched intestinal organoids and tumouroids. Nat. Biomed. Eng. 8, 345–360 (2024).

84. Recaldin, T. et al. Human organoids with an autologous tissue-resident immune compartment. Nature 633, 165–173 (2024).

85. Lancaster, M. A. & Knoblich, J. A. Generation of cerebral organoids from human pluripotent stem cells. Nat. Protoc. 9, 2329–2340 (2014).

86. Bai, H. et al. Mechanical control of innate immune responses against viral infection revealed in a human lung alveolus chip. Nat. Commun. 13, 1928 (2022).

87. Dasgupta, Q. et al. A human lung alveolus-on-a-chip model of acute radiation-induced lung injury. Nat. Commun. 14, 6506 (2023).

88. Völkner, M. et al. Retinal Organoids from Pluripotent Stem Cells Efficiently Recapitulate Retinogenesis. Stem Cell Rep. 6, 525–538 (2016).

89. Hötzel, K. J. et al. Synthetic Antigen Gels as Practical Controls for Standardized and Quantitative Immunohistochemistry. J. Histochem. Cytochem. 67, 309–334 (2019).

